# A whole-brain imaging-based systems approach to understand origin of addiction in binge-like drinking model

**DOI:** 10.1101/2021.02.17.431586

**Authors:** Marzena Stefaniuk, Monika Pawłowska, Klaudia Nowicka, Marcin Barański, Zbigniew Zielinski, Łukasz Bijoch, Diana Legutko, Piotr Majka, Sylwia Bednarek, Natalia Jermakow, Daniel Wójcik, Leszek Kaczmarek

## Abstract

Many fundamental questions on addiction development are still unanswered. These questions are frequently difficult to address by examining a single brain structure, but can best be addressed at the systems level. Neurons create functional networks that change over time, since brain regions may work together differently in different contexts. We offer a framework for describing the nature behind alcohol binge drinking and the transition to addiction. The present study investigated whole-brain c-Fos expression following reexposure to alcohol in a model of binge-like drinking in mice in IntelliCage. We developed a dedicated image computational workflow to identify c-Fos-positive cells in three-dimensional images obtained after optical tissue clearing and whole-brain imaging in the light-sheet microscope. We analyzed functional networks and brain modularity following reexposure to alcohol. c-Fos levels in brains from animals that were reexposed to alcohol were clearly different from binge drinking animals. Structures involved in reward processing, decision making and characteristic for addictive behaviors stood out particularly. In alcohol reexposed animals differently active structures either gained or lost correlation when compared to the control group.

## Introduction

Binge drinking is a type of alcohol use disorder (AUD) that involves consuming an excessive amount of alcohol in a short time (2004). Binge drinking episodes may turn into heavy drinking and increase the likelihood of developing addiction (Dawson *et al*, 2005). In alcohol addiction, cues that are associated with alcohol trigger craving and the relapse (George & Koob, 2010; Namba *et al*, 2018). Alcohol induces multiple alterations in the brain that are thought to contribute to addiction, but an understanding of the neuronal ensembles responsible for excessive alcohol drinking is still preliminary (George & Hope, 2017).

Whole-brain mapping techniques produce large datasets of anatomical and functional connectivity patterns (Sporns & Betzel, 2016). The state of functional brain networks may influence the ways in which the brain manages information and changes in result of experience (Rubinov & Sporns, 2010). Strong evidence shows that brain networks have a modular organization [for review, see (Bullmore & Sporns, 2009)]. Such modules are thought to represent groups of brain regions that are collectively involved in specific functional domains (e.g., supporting cognition) (Sporns & Betzel, 2016). Brain network properties, such as modularity, provide an important dimension for understanding the brain in health and disease (Arnemann *et al*, 2015; Liu *et al*, 2019; Meunier *et al*, 2009; Zhang & Volkow, 2019).

Whole-brain imaging in human subjects has advanced our understanding of the effects of alcohol on the brain, but its use has been limited by ethical considerations, the heterogeneity of alcoholic populations, and the low spatial resolution of imaging methods (Zahr & Pfefferbaum, 2017). In contrast, animal models enable the control of multiple genetic, environmental, and alcohol consumption factors (Fuchs *et al*, 2019; Spanagel, 2017). By mimicking neuropsychiatric criteria, such models help mimic patterns observed in humans (Sanchis-Segura & Spanagel, 2006; Skora *et al*, 2020). Furthermore, spatial resolution of the imaging approaches that are available for studying animals’ brains are far superior over those available in humans (Martin, 2014). Identifying neurons, their pathways and networks that modulate binge drinking may provide insights into what happens when a subject is occasionally exposed to alcohol. Thus, this knowledge may eventually lead to developing therapies against addiction.

The present study sought to provide a global analysis of the pattern of binge-like drinking-related c-Fos expression in the brain in mice. c-Fos is a widely used marker of neuronal activation that mediates the long-term potentiation of excitatory synapses (Knapska *et al*, 2007; Sun *et al*, 2020). We adapted the drinking-in-the-dark (DID) binge-drinking paradigm (Rhodes *et al*, 2005; Thiele & Navarro, 2014) to the IntelliCage training system. This behavioral setup allows the long-term monitoring of animals that live in social groups with minimal contact with experimenters (Beroun *et al*, 2018; Kiryk *et al*, 2020; Koskela *et al*, 2018; Radwanska & Kaczmarek, 2012; Smutek *et al*, 2014; Stefaniuk *et al*, 2017). Training consisted of initial acquaintance with alcohol that was presented concomitantly with a visual cue, followed by a withdrawal period and next by a session that exposed the mice to either the cue alone or the cue with alcohol access, as during the initial training. To assess c-Fos expression that was triggered by reexposure, the brain tissue was collected 2 h after alcohol administration. Importantly, c-Fos expression typically follows a very specific time course, in which protein is detectable in neuronal nuclei within ~1-2 h after a stimulation (Knapska *et al.*, 2007). To reveal brain-wide neuronal activation patterns, whole brain hemispheres were optically cleared and immunostained for c-Fos. Clearing removes lipids to render the tissue transparent (Chung *et al*, 2013; Renier *et al*, 2014; Susaki *et al*, 2015). Cleared hemisphere were imaged *in toto* using a light-sheet microscope (Pawlowska *et al*, 2019). The signal intensity of c-Fos-positive nuclei was transformed into the signal density that was specific to each brain structure. These were next used to examine whether reexposure to alcohol or the alcohol-associated cue following long-term withdrawal resulted in alterations of c-Fos expression, functional networks, and brain modularity.

## Results

### Adaptation of the drinking-in-the-dark model to the IntelliCage training system

We employed an animal model of binge-like alcohol drinking, referred to as DID (Thiele & Navarro, 2014) that was adapted for the IntelliCage (Fig. 1A). This behavioral training system allows the tracking of corner visits, nosepokes that permitted access to and licks from the drinking bottles located in the corners (Fig. 1B). Only one mouse could be present in a corner at a time. Previously, the usefulness of the IntelliCages for modeling AUD has been demonstrated (Kiryk *et al.*, 2020; Radwanska & Kaczmarek, 2012; Stefaniuk *et al.*, 2017).

**Figure 1.**
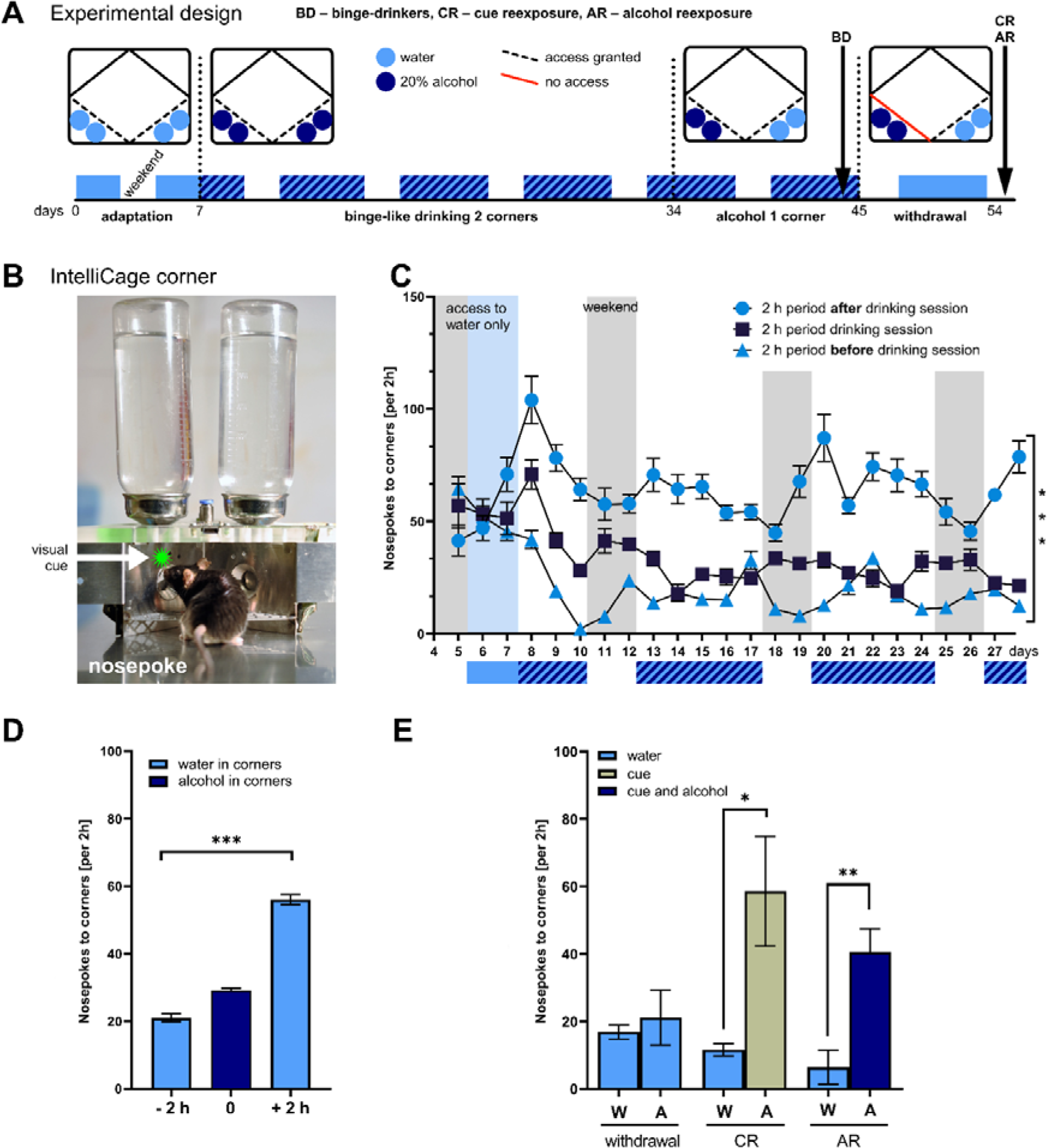
Binge-like alcohol drinking in the IntelliCage. (A) Experimental design and cage configuration. Blue circle: access to water. Dark blue circle: access to alcohol. Dashed line: free access. Solid red line: no access. (B) IntelliCage corner. The mice had access to two corners that were equipped with a door that controlled access to the bottles. Whenever access to alcohol was granted, a visual cue (a green LED over the door) turned on. (C) Nosepokes in the corners at 2-h intervals. The mice actively sought alcohol when no access to alcohol was granted (nosepokes in the corners, two-way ANOVA, *F*_2,1005_ = 511, \*\*\**p* < 0.0001). For each day, three time intervals are presented: blue triangle (2-h period before alcohol access), dark blue square (2 h of alcohol access in DID session), blue circle (2-h period after 2 h of alcohol access). The data are expressed as mean ± SEM. (D) Increase in the number of nosepokes in corners when access to alcohol was ceased (two-way ANOVA, *F*_2,945_ = 367.8, \*\*\**p* < 0.0001). (E) Behavioral phenotyping upon reexposure. The mice made more nosepokes in the corner with alcohol following reexposure compared with the water corner in CR group *p* = 0.0206, *t*_8_ = 2,878 and AR group *p* = 0.0015, *t*_8_ = 4,746. W, water corner; A, alcohol corner (during withdrawal, A indicates the corner with the previous alcohol access); CR, cue reexposure; AR, alcohol reexposure. The data are expressed as mean ± SEM. \**p* < 0.05, \*\**p* < 0.01 (*t*-test).

In the original DID design that was reviewed by Thiele and Navarro (Thiele & Navarro, 2014), mice were housed individually. Three hours into the dark phase of the diurnal cycle, instead of water, they were given access to 20% alcohol for 2 h. The procedure lasted for 4 days. At the end of each drinking session, blood samples were collected to measure blood alcohol levels (Cox *et al*, 2013).

In the present experimental design, we followed the same timeframe of drinking sessions (2 h session starting 3 h into the dark phase of the light/dark cycle), but the mice were kept in groups and did not undergo the sessions individually. Five seconds of access to alcohol was provided after each nosepoke, which was always associated with a visual cue (Fig. 1B). We extended the DID sessions to 3 weeks (with no access to alcohol during weekends as per original protocol). For the next 2 weeks, we limited access to alcohol to one corner with concurrent access to water in another corner. It was followed by a 9-day withdrawal with no access to alcohol. Finally, the mice were exposed to either 2-h access to alcohol that was paired with a light cue or the cue only in the corner that was previously associated with alcohol (Fig. 1A). Importantly, we did not attempt to measure blood alcohol levels to exclude possible confounding effects of manipulation, such as stressing the animals by separating them from the group.

### Mice exhibit an increase in craving for alcohol after drinking sessions

During pre-binge drinking period, the mice explored the cage and had free access to drinking water in two corners. They learned to make a nosepoke to access the drinking bottles. Next, the mice were given access to 20% alcohol, combined with a visual cue, that was provided in the corners. During each drinking session, we measured the number of alcohol licks made by each mouse. This measure was used as a proxy of alcohol consumption. Throughout the experiment, the animals consumed alcohol voluntarily during daily 2-h sessions, reaching an average consumption of 3.82 ± 0.88 mg/kg of body weight per 2 h per day, when alcohol was available in two corners, and 3.57 ± 1.23 mg/kg of body weight when alcohol was limited to one corner. With our DID protocol, the animals developed craving for alcohol. This was manifested by a marked increase in the number of nosepokes in the alcohol corners during the 2-h period immediately following the drinking session (Fig. 1C, D). Throughout the drinking sessions, the mice made an average of 16 ± 3 nosepokes in the corners. During the 2 h immediately before the drinking sessions, they made on average of 10 ± 2 nosepokes in the corners. Notably, 2 h after the drinking session, they made an average of 33 ± 7 nosepokes in the corners, indicating high levels of craving for alcohol. The number of nosepokes between these three time periods was significantly different (Fig. 1C). We observed a similar result when we analyzed the average number of nosepokes per session (Fig. 1D). During weekends, when alcohol was not provided, these seeking patterns were also maintained. These results indicate that the animals actively sought alcohol, predominantly immediately after binge-like drinking sessions.

### Mice prefer alcohol over water during reexposure to alcohol or the alcohol-related cue

After the drinking period, the animals were deprived of access to alcohol for 9 days, followed by reexposure. To verify that behavior during reexposure was unrelated to overall activity, we performed an analysis of the correlation between the number of nosepokes during this period and the number of all nosepokes in all corners during the entire experiment and found no correlation (Supplementary Fig. S1).

Under reexposure conditions, we measured the number of nosepokes in the alcohol corner and water corner and compared them to the corresponding timeframe during the withdrawal session that considered activity during the last day of withdrawal. In the CR group, a cue (light) appeared following a nosepoke, but no alcohol reinforcement was provided. In the AR group, each nosepoke opened the door to grant access to alcohol for 5 s simultaneously with the visual cue. Animals in both groups preferred the alcohol corner over the water corner (average of 59 ± 36 and 12 ± 4 nosepokes in the alcohol corner and water corner, respectively, in the CR group; average of 41 ± 15 and 6 ± 5 nosepokes in the alcohol corner and water corner, respectively, in the AR group, Fig. 1E). These results indicate that mice that were exposed to alcohol and the alcohol-associated cue in the binge-like drinking model exhibited a preference for the alcohol corner, suggesting that they associated the alcohol corner and the alcohol-related cue with alcohol reward.

### Whole-hemisphere c-Fos expression profiling in binge-like drinking mice

To examine whether reexposure to alcohol or the alcohol-related cue following withdrawal after prolonged binge-like drinking resulted in changes in c-Fos expression, we performed optical tissue clearing and c-Fos immunohistochemistry (Renier *et al.*, 2014). To prepare a map of brain areas that responded to either alcohol plus the cue or the cue alone, we aligned the brain structure distribution of the c-Fos signal with the *Allen Brain Atlas* (2017). The overall result was a three-dimensional map of c-Fos-positive puncta with specific anatomical annotations (see Supplementary Movie S1). This map was converted to a c-Fos signal density map that was specific for each animal. A hierarchical list of identified structures was used for comparisons between groups, correlation analysis, structural modularity analysis and network analysis.

Several brain structures differentially expressed c-Fos in response to specific experimental conditions (Supplementary Dataset S1). These included structures that are implicated in motor function, visual and olfactory areas, the thalamus, and other. The most prominent differences were found between the AR group and BD and CTRL groups. The c-Fos signal density was much higher in the emotional response-related infralimbic area (ILA), decision-making-related anterior cingulate (ACA), motor function-related cuneiform nucleus (CUN), motivation process-related pericommissural nucleus (PRC), and memory-related postrhinal area (VISpor) in the AR group compared with the CTRL and BD animals. For statistical details for other structures, see Supplementary Dataset S1 and Fig. S2-S5.

### Alcohol reexposure but not binge-like drinking induces activation of the amygdala

c-Fos expression in the amygdala in rats was previously found to be markedly elevated following reexposure to alcohol-related cues (Radwanska *et al*, 2008). In the present study, we noted a clear difference in c-Fos expression in the AR group, particularly in amygdala nuclei and their afferent and efferent structures. The central nucleus of the amygdala (CEA), basolateral amygdala anterior part (BLAa) exhibited a higher c-Fos signal density in the AR group compared with the BD group (Fig. 2). Another structure that was prominently activated by alcohol reexposure was the bed nucleus of stria terminalis (BST), a major output pathway of the amygdala, and the ventral tegmental area (VTA), an amygdala input. One of the richest areas of c-Fos-positive nuclei in the AR group was the anterior cingulate area (ACA), a structure that provides a strong input to the BLA. For detailed comparisons between the BD and AR groups, see Fig. 3. For comparisons between the other groups, see the Supplemental Information. These results indicate that during reexposure to alcohol after withdrawal, the whole amygdala complex and its input and output structures are strongly activated, reflected by c-Fos expression, compared with BD animals.

**Figure 2.**
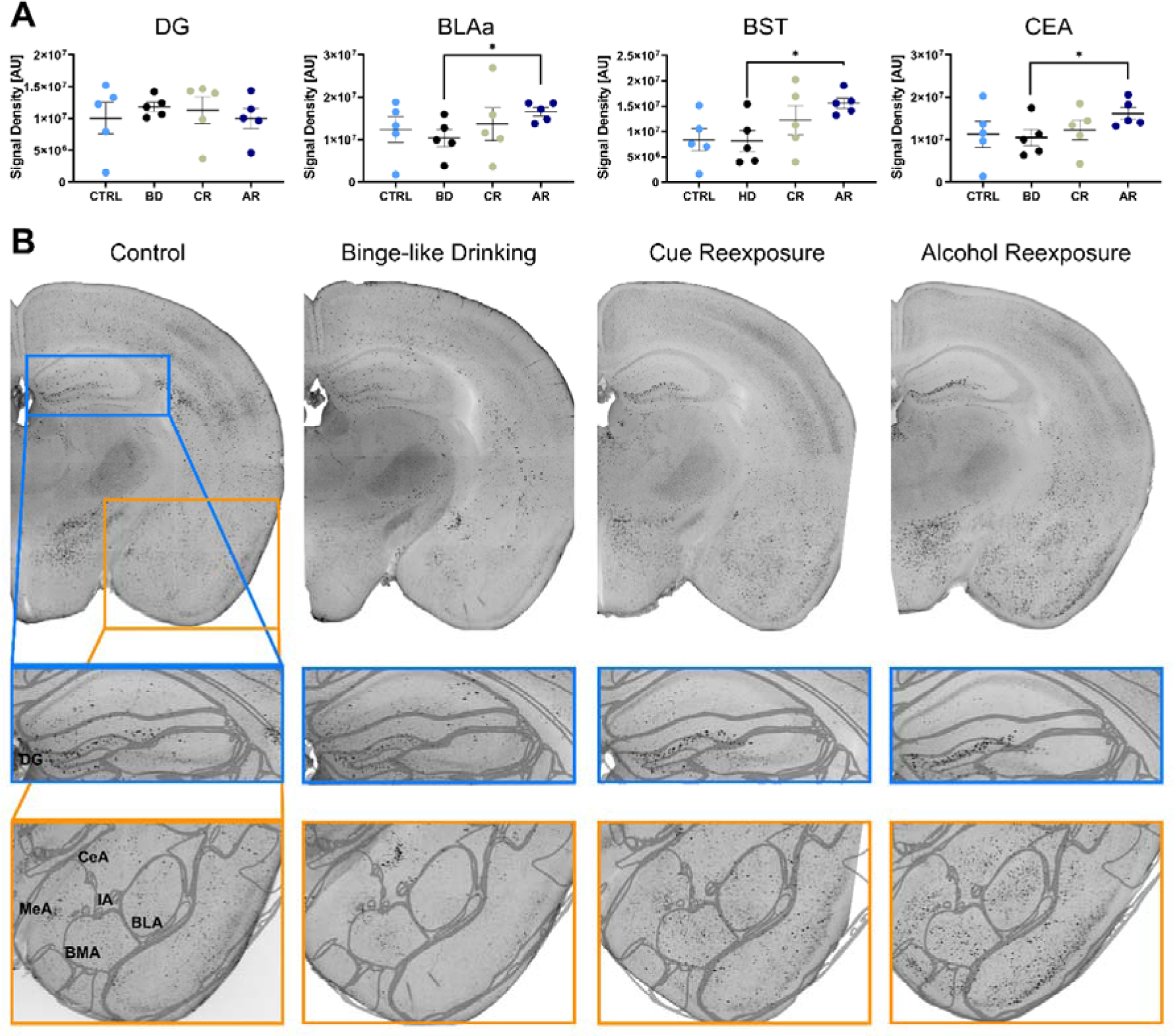
Overview of profile of c-Fos expression in selected brain structures. (A) The signal density for each structure was obtained by dividing the sum of the signals of individual c-Fos-positive cells in a given structure by its volume (in mm^3^). \**p* < 0.05, \*\**p* < 0.01, \*\*\**p* < 0.001 (*t*-test). No difference in the c-Fos signal density was found in the dentate gyrus between experimental groups and increase in signal density in selected amygdala nuclei and BST was observed. (B) Coronal sections of maximum intensity projections that covered 50 μm, reconstructed from whole-brain light-sheet microscopy images. Single c-Fos-positive nuclei in specific brain areas and differences between experimental groups are shown. BLA, basolateral amygdala; BST, bed nucleus of stria terminalis; CeA, central nucleus of the amygdala; DG, dentate gyrus; MeA, medial amygdala; Control, control naive group with access to water only; Binge-like Drinking group with no history of alcohol withdrawal; Cue Reexposure, alcohol-related cue reexposure group; Alcohol Reexposure, alcohol reexposure group.

**Figure 3.**
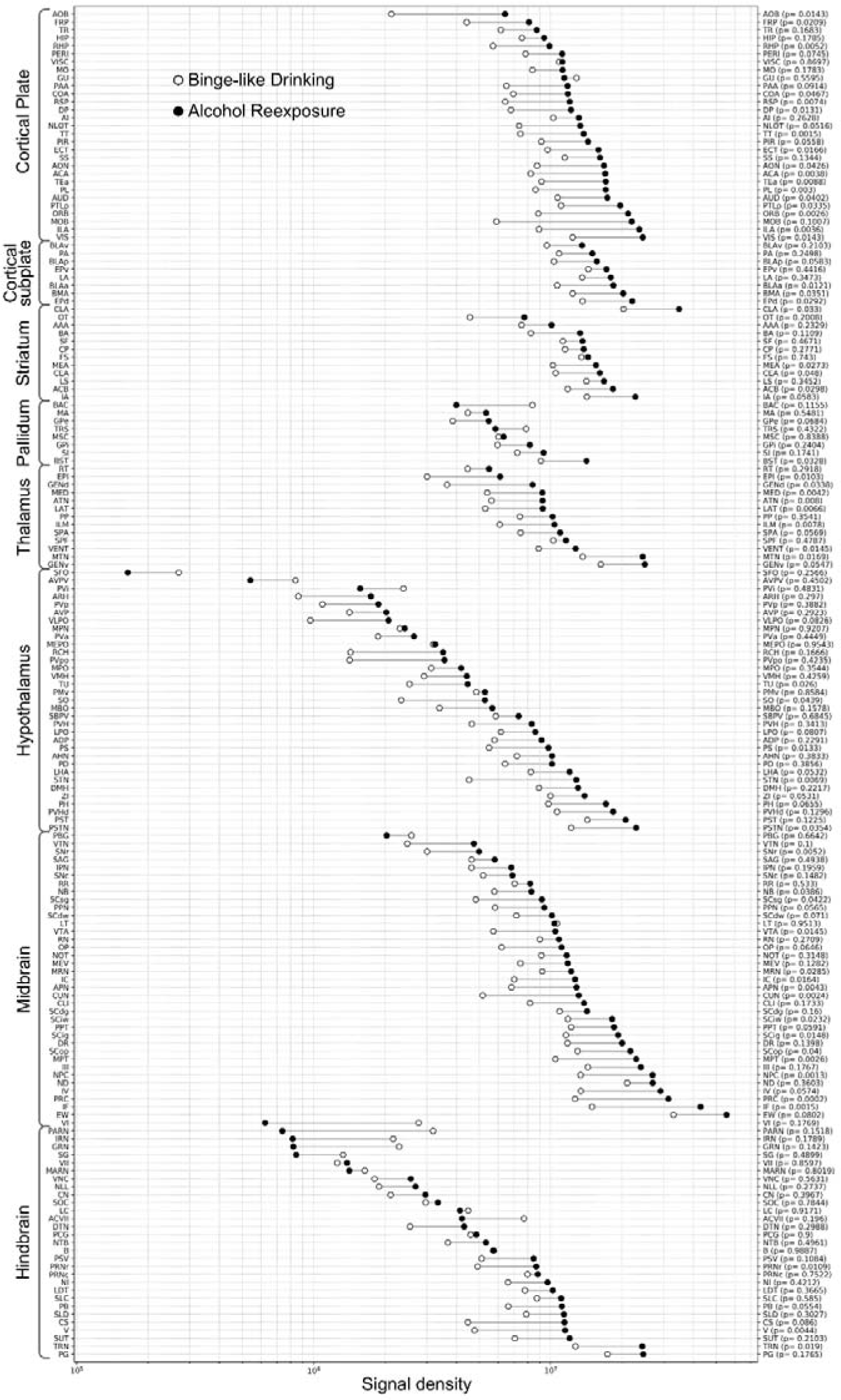
Overview of the c-Fos signal density profile in the whole brain, showing a comparison between the binge-like drinking group and alcohol reexposure group. The signal density for each structure was obtained by dividing the sum of signals of individual c-Fos-positive cells in a given structure by its volume (in mm^3^). The structures are grouped in anatomical categories according to the *Allen Brain Atlas*. Within-category structures are shown according to decreasing signal density. The data were analyzed using one-way paired *t*-test, and resulting *p* values are in brackets next to the structure acronym.

### Reexposure to alcohol alters functional neuronal co-activation

To examine whether reexposure to alcohol or the alcohol-related cue resulted in functional network alterations, we used data on c-Fos profiling and performed structural co-activation analysis by calculating Pearson correlation matrices for each group separately. The ordering of structures in these matrices followed the anatomical groups from the *Allen Brain Atlas* (2017). The following gross divisions of the brain were applied: cortical plate, cortical subplate, striatum, pallidum, thalamus, hypothalamus, midbrain, and hindbrain. We observed similar patterns of coactivation in the CTRL and CR groups (Fig. 4A, C). Binge-like drinking mice exhibited a lower level of cross-correlation between various brain regions compared with the CTRL group (Fig. 4B). The AR group exhibited a prominent anti-correlation between more brain regions compared with all of the other groups, with the exception of hypothalamus nuclei, which had a strong interregional positive correlation (Fig. 4D). Reexposure to alcohol was shown to activate the amygdala; thus, we analyzed functional connectivity of amygdala nuclei with other brain regions and found that the amygdala became either poorly or negatively correlated in the AR group (Fig. 4E). For details see the Supplement. These results indicate that the activity of brain regions in response to alcohol was less correlated globally compared with alcohol drinking, binge drinking, or exposure only to the alcohol-related cue. Reexposure to alcohol induced the most prominent and distinctive brain-wide changes.

**Figure 4.**
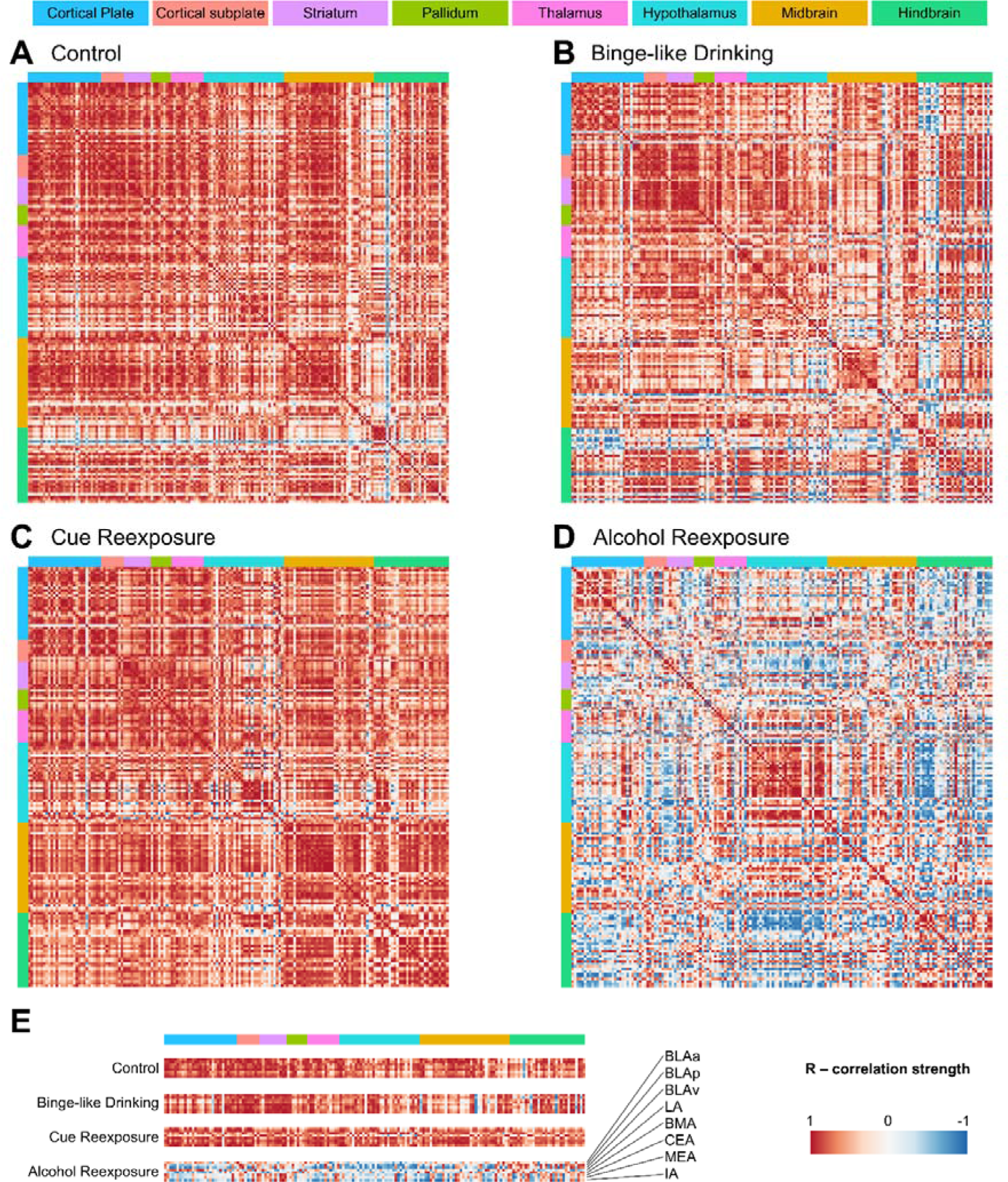
Brain region correlation heatmaps organized according to the *Allen Brain Atlas*. (A) Correlation map for the naive control group that had access to water only and no previous exposure to alcohol. (B) Correlation map for the binge drinking group whose brains were snap frozen after the last drinking session. (C) Correlation map for the cue-reexposed group following alcohol withdrawal. (D) Correlation map for the alcohol reexposure group following alcohol withdrawal. Color-coded anatomical annotations are depicted above the heatmaps. R indicates the correlation strength between single structures, in which warmer colors indicate more similar signal densities between two structures in all mice (*n* = 5) in a given group. Cooler colors indicate a lower correlation. (E) Comparison of correlations of amygdala structures *vs*. all other regions.

### Reexposure to alcohol increases structural modularity of the brain

Brain networks are information processing systems that share a modular community structure (Meunier *et al*, 2010). A network is modular when sets of structures form densely interconnected groups with sparser connectivity between groups (Newman, 2006). Brain modularity alterations have been described in numerous brain disorders (Arnemann *et al.*, 2015; Liu *et al.*, 2019) and also upon withdrawal from alcohol consumption (Kimbrough *et al*, 2020). To examine whether reexposure to alcohol or the alcohol-related cue following withdrawal after prolonged binge-like alcohol drinking resulted in changes in brain modularity, we performed hierarchical organization analysis (Fig. 5). We calculated the Euclidean distance between rows of the correlation matrix and performed hierarchical clustering (Fig. 5). Heatmaps were generated with dendrograms in which the height at which any two structures were first connected corresponded to their degree of similarity. To compare dendrograms of the different experimental groups, we plotted the number of clusters based on relative tree heights. The hierarchical cluster dendrograms were next cut at half the height of each given tree to split the dendrogram into specific modules. In control animals, the analysis revealed that the brain was organized into three modules (Fig. 5A). In mice that were exposed to alcohol, the brain was organized into seven modules in the BD group (Fig. 5B), eight modules in the CR group (Fig. 5C), and 12 modules in the AR group (Fig. 5D). This indicates that binge-like drinking itself increased brain modularity. Reexposure to alcohol following withdrawal increased hierarchical modularity of the brain even further (Fig. 5E).

**Figure 5.**
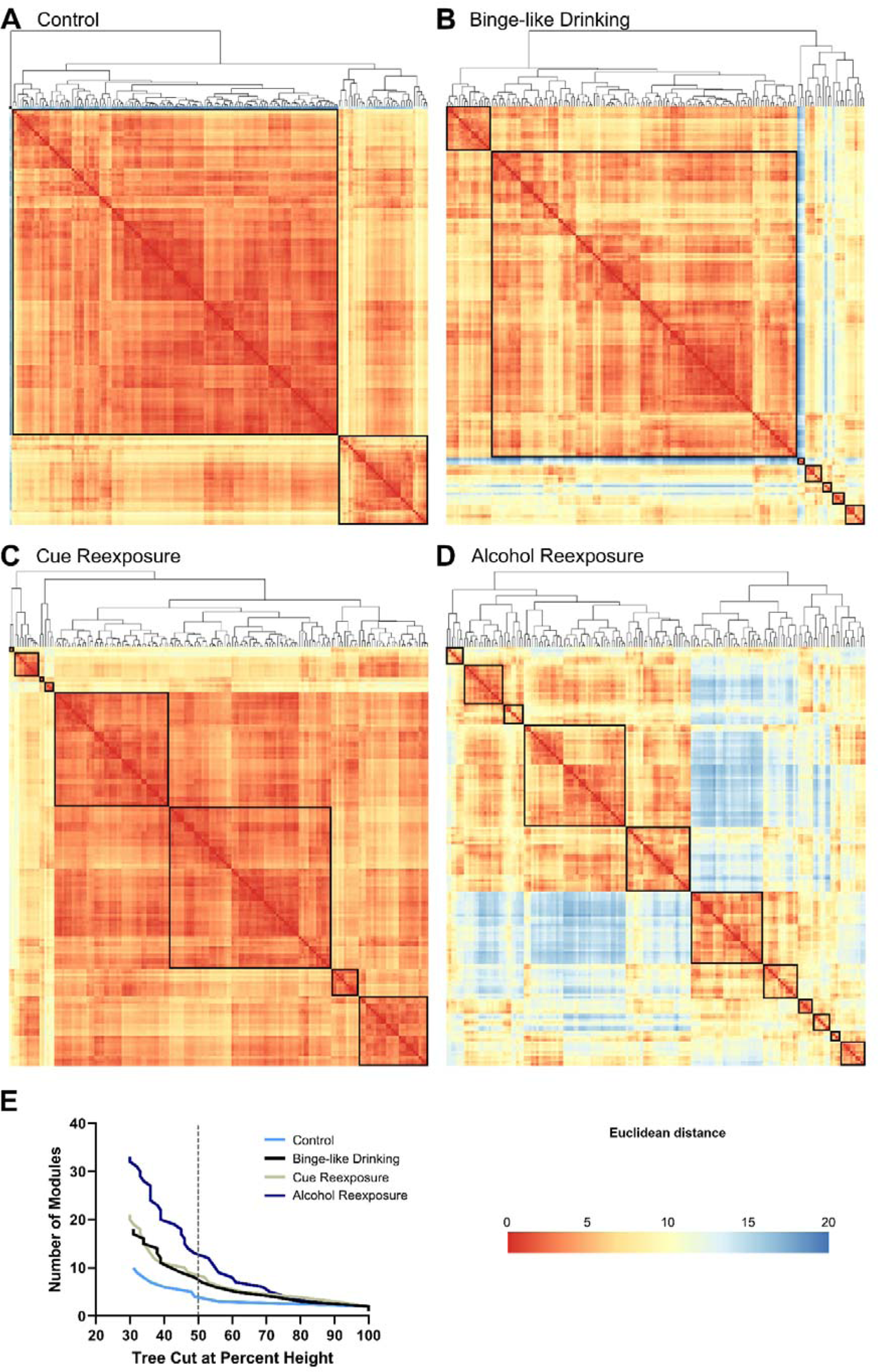
Intra-network hierarchical organization of sub-networks, showing the relative distance between brain regions in the (A) control group, (B) binge-like drinking group, (C) cue reexposure group, and (D) alcohol reexposure group. The Euclidean distance was calculated to create the dendrograms. The lines in the dendrograms are drawn in proportion to the distance measure of the nodes. The sub-networks are in ascending order according to their distances in the sub-dendrogram. Warmer colors indicate a shorter Euclidean distance, indicating that two structures have highly similar c-Fos signal density correlation patterns with all other structures. Black squares indicate modules that were created by cutting the dendrograms at half the maximal tree height. (E) Number of modules per group based on cutting the dendrograms at different percentages of tree heights. The dashed line indicates cutting at 50% to create modules in A-D.

We also found that the amygdala and its input and output structures that were shown to be active with reexposure to alcohol shared one module in CTRL, shared two modules in BD animals, and fell into different modules in AR animals, thus indicating a major network rearrangement (Supplementary Dataset 2 and Fig. S11-14).

Finally, we investigated whether we could distinguish any specific functional module with components that are involved in substance use disorder or reward circuitry in the AR group (Koob & Volkow, 2016). We identified two modules that included five structures that are implicated in addiction (i.e., ventral tegmental nucleus [VTN], temporal association area [TEa], BST, superior colliculus [SCiw], and nucleus accumbens [ACB]). Interestingly, when comparing c-Fos density profiles in the AR group, the TEa, BST, SCiw, and ACB had a higher level of c-Fos accumulation compared with binge-like drinkers (see Fig. 3). We also identified another potentially relevant module whose structures are implicated in reward-seeking behavior. This module included the substantia innominata (SI), retrohippocampal region (RHP), interfascicular raphe nucleus (IF), and BLAa. The BLAa is one of the major components in this module, together with the IF and subiculum (part of the RHP). Other amygdala components shared one module (medial amygdala [MEA], subparafascicular nucleus [SPF], vestibular nuclei [VNC], CEA, postpiriform transition area [TR], parastrial nucleus [PS], red nucleus [RN], intercalated nuclei [IA], pedunculopontine nucleus [PPN], supratrigeminal nucleus [SUT], subfornical organ [SFO], ectorhinal [ECT], and visceral area [VISC]).

These results indicate that alcohol reexposure implicates disconnection of functional networks of the brain toward the formation of fine circuitries that are involved in the development of addiction and processing of emotional information. Importantly, no such rearrangements were observed in the control group or binge-like drinking animals after the last drinking session.

### Network fragmentation in alcohol reexposed brain

To assess whether modular organization of the cue and alcohol reexposed brain network was related to neurobiological organization of the structures relevant to addiction (Koob & Volkow, 2016), we employed a functional network analysis, as a collection of nodes (structures) and edges (links) between pairs of nodes (Rubinov & Sporns, 2010). Individual nodes and edges differ in their impact on the overall functioning of the network. Some nodes are more essential or more influential than others. Some edges carry more traffic. Important nodes are often more highly and densely connected to the rest of the network facilitating global integrative process (Sporns, 2010). First, to identify interaction between all structures we calculated within-module degree z-score (WMD) and the participation coefficient (PC). The role of each node was defined by both of those values (Guimera & Nunes Amaral, 2005) (Fig. S15), where WMD sets the importance of structure within its module and PC the importance of outside module connections. All nodes were classified into four distinct groups: provincial hubs (likely to play an important part in the facilitation of modular segregation), connector hubs (highly relevant outside and inside module), non-hub connectors (likely to facilitate global intermodular integration) and peripherals (with no role in within or outside connectivity) (for details see Supplementary Methods). We noticed that in reexposure groups some structure gain functionality facilitating intermodular integration (Fig. S15).

Next, we selected brain structures relevant to addiction development: ACB, AI, BLA (divided into BLAa, BLAp, BLAv), BST, CEA, GPe, GPi, HIP, IA, SNc, VTA (Koob & Volkow, 2016) and performed functional connectivity analysis (Fig. 6). In animals living in homecages with no access to alcohol, spontaneous “resting-state” functional connectivity state revealed most structures were equally connected with similar strength (Fig. 6A). After two-hour drinking session a pattern of large functional connectivity network changed with Globus Pallidus segments being slightly detached (Fig. 6B). While exposure to alcohol matched cue following withdrawal did not significantly changed the network (Fig. 6C), alcohol reexposure following withdrawal caused a major disconnection of functional network of the brain (Fig. 6D). Reexposure to alcohol implicated the disruption of functional interactions between structures creating specific patterns where BST and VTA shared the same pattern and it is different from GPe, BLAp, BLAv, HIP, IA, SNc and CEA pattern (Fig. 6D). We found that the extended amygdala structure CEA and BST and cortico-striatal loops still interacted with the mesolimbic dopamine region, however, the relationship is much weaker with almost no interactions with structures within the region. Interestingly ACB did not interact through positive connectivity with any other addiction relevant structure (Fig. 6D).

**Figure 6.**
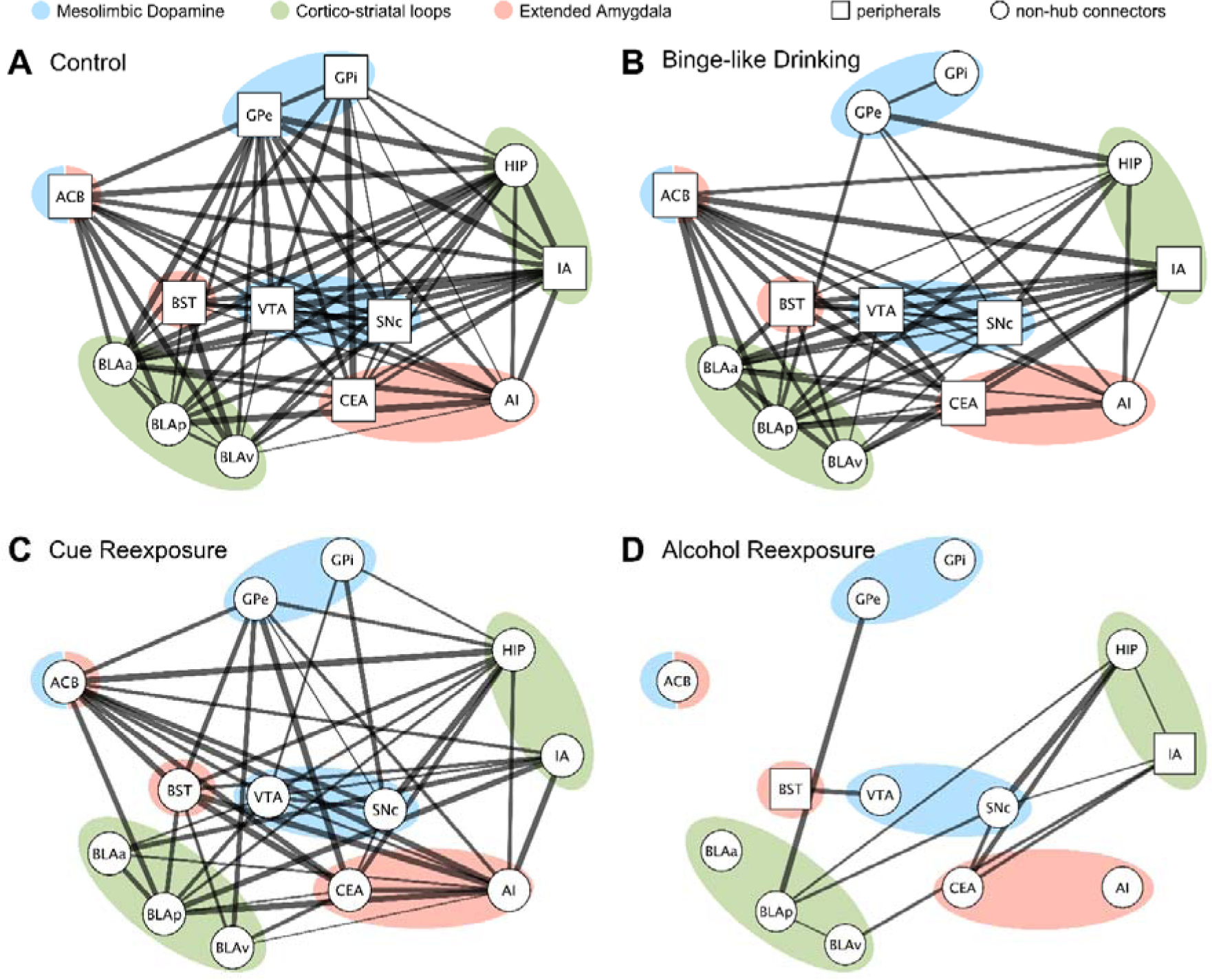
Functional connectivity of brain regions relevant to addiction. Each node (square or circle) represents a structure. The shape of the node represents the importance of node within module and between modules (see Supplementary Figure) with non-hub connectors (circles) being important connecting different modules and peripherals having equal role within and outside module. The thickness of the lines (edges) represents the strength of the correlation, with thicker line representing higher correlation. Gross anatomy is represented by colored bubble areas. Note loss of edges in binge-like drinking animals (B) and further in cue (C) and alcohol reexposed animals (D) when compared to control animals (A) indicating alterations of functional interactions from disruption to strengthening. Structures selected from (Koob & Volkow, 2016). For abbreviations see Supplementary Dataset 1.

## Discussion

In this study, we characterized global brain c-Fos activity in a mouse model of binge-like alcohol drinking. First, we evaluated binge-like drinking behavior in IntelliCage. Next, we identified brain regions that exhibited alterations of c-Fos signal density following reexposure to alcohol, using whole-brain imaging technique on optically cleared tissue. We observed different brain structure correlation patterns in naive animals and alcohol- or cue-reexposed animals. Lastly, we observed an increase in brain modularity and decrease in functional connectivity in animals that were reexposed to alcohol, compared with all of the other groups.

To characterize the binge-like drinking phenotype in mice, we adapted the DID model (Rhodes *et al.*, 2005; Thiele & Navarro, 2014) to the IntelliCage system. This behavioral training and testing system was previously used to model alcohol addiction in mice (Koskela *et al.*, 2018; Radwanska & Kaczmarek, 2012; Smutek *et al.*, 2014; Stefaniuk *et al.*, 2017). To date, however, no study has used the IntelliCage to study binge-like alcohol drinking.

The DID assay provides a 20% alcohol solution to mice for a limited time each day. In the original protocol, exposure to alcohol was limited to 2 h, 3 h into the dark phase of the diurnal cycle, for 4 days, accompanied by daily measurements of blood alcohol levels. In the current experiment, we extended the procedure and omitted blood alcohol level measurements to reduce stress in the animals. The use of IntelliCage allowed tracking individual mice movement and calculating alcohol consumption with high accuracy, that reached levels consistent with experiments which used standard DID method (Rhodes *et al*, 2007; Wilcox *et al*, 2014). Even with concurrent access to water in the last DID session, the animals still reached similar levels of alcohol consumption. A similar observation was made in a previous study (Giardino & Ryabinin, 2013).

Using the IntelliCage DID training paradigm, we found an increase in alcohol craving immediately after binge drinking session. The persistence of seeking alcohol has been previously used in other models of AUD as a measure of craving (Giuliano *et al*, 2018; Radwanska & Kaczmarek, 2012; Stefaniuk *et al.*, 2017); for review, see (Luscher *et al*, 2020). Furthermore, the mice exhibited an increase in preference for either alcohol or the cue alone following reexposure after withdrawal. Similarly, previous studies showed that a history of alcohol drinking during DID increased alcohol preference (Cox *et al.*, 2013; Wilcox *et al.*, 2014). Considering the complexity of AUD and the number of protocols that are used to model it based on various aspects of drinking patterns, our approach may facilitate further studies of the transition to addiction in automated controlled systems, such as the IntelliCage.

To create a map of brain structures that responded to either alcohol or the cue following withdrawal, we used c-Fos expression as a proxy of neuronal activation. c-Fos is a transcriptional factor that is activated upon neuronal stimulation (Knapska *et al.*, 2007; Sun *et al.*, 2020; Yap *et al*, 2020). Transcriptional modifications are postulated to be crucial for the development of addictive states (Nestler, 2001). Alterations of gene expression would lead to lasting modifications of neuronal activity and ultimately changes in neural circuits and stable changes in behavior. c-Fos has been reported to be activated after reexposure to alcohol (Smith *et al*, 2019) and alcohol-related cues (Radwanska *et al.*, 2008). The cue- and context-induced reinstatement of seeking alcohol induces the expression of c-Fos in discrete populations of neurons in various brain regions (Hamlin *et al*, 2009; Hamlin *et al*, 2007; Marinelli *et al*, 2007; Millan *et al*, 2010). Nonetheless, an overall picture of c-Fos expression is still lacking. In the present study, to trigger c-Fos expression, the mice were exposed to either alcohol or the cue after a 9-day withdrawal. Our analysis revealed a complete map of structures that were characterized by different activity profiles, reflected by the density of c-Fos expression. A similar approach was used after antipsychotic treatment with haloperidol (Renier *et al*, 2016), the regulation of energy expenditure (Schneeberger *et al*, 2019), alcohol abstinence (Kimbrough *et al.*, 2020) and opioid relapse (Keyes *et al*, 2020).

The present study advances the c-Fos mapping technique by introducing c-Fos signal density analysis with more accurate anatomical details that cover many structures and layers. For instance, we identified various densities of c-Fos signals in different layers and parts of the ACA in animals that were reexposed to alcohol. Noteworthy, ACA in humans has been reported to contribute to motivational valence assignment (Mesulam, 1990) and reward assessment (Knutson *et al*, 2000). Other structures that are markedly activated in animals that are reexposed to alcohol include nuclei of the amygdala that have been previously implicated in reinforcing drug craving (Ryabinin *et al*, 1997) and addiction in studies in human patients (Goldstein & Volkow, 2002; Koob & Volkow, 2016) and animal models (Augier *et al*, 2018). Markedly altered were also input and output structures of amygdala nuclei implicated in drug seeking (Belin *et al*, 2013). The global brain c-Fos profiling approach provides information about particular structures and their neuronal ensembles. Such profile of neuronal activation may shed light on addiction with regard to notable brain regions and specific patterns of sparsely distributed neurons (i.e., neuronal ensembles) that are selectively activated by specific cues, contexts, and rewards during alcohol-related behavior (George & Hope, 2017).

Further analysis of c-Fos density correlation maps enabled comparisons between groups of structures. All such groups exhibited specific correlation patterns, with high positive correlations in control binge-like drinking animals and cue-reexposed animals. Following reexposure to alcohol, groups of structures were highly negatively correlated. This indicates a significant reconstruction of the global brain activity pattern. A recent study showed that such rebuilding is also observed during abstinence, although in the opposite direction (i.e., positive co-activation) (Kimbrough *et al.*, 2020). In the present study, however, the trigger of c-Fos expression was different and had a well-defined onset (i.e., reexposure to alcohol).

Using hierarchical clustering, we found an increase in brain modularity in the AR group (12 functional networks/modules) compared with the BD, CR, and CTRL groups showing seven, eight and three modules respectively. Changes in brain modularity have also been observed during abstinence (Kimbrough *et al.*, 2020), with a decrease in brain modularity. We hypothesize that reexposure to alcohol can be a strong behavioral stimulus that is sufficient to trigger major changes in functional brain architecture. Kimbrough *et al.* (Kimbrough *et al.*, 2020) reported that a decrease in modularity might be partially responsible for cognitive dysfunction seen in humans and animal models of alcohol dependence, but this does not appear to be the case in binge-like drinking models. Human magnetic resonance imaging studies indicate that network organization into higher modularity and thus connectivity within modules and sparse connections between modules may reflect more effective signaling across brain regions (Stevens *et al*, 2012). We suggest that the increase in brain modularity that was observed under the present experimental conditions might be transient and limited selectively to reexposure to alcohol and not to alcohol consumption itself, in which BD animals had lower modularity than the AR group.

Finally, we aimed at connecting practically all structures into interacting network defining function of components within the network (Sporns, 2010). We described the effective strength of functional interaction and contributions of individual nodes determined by their interaction within a local and global neighborhood (Guimera & Nunes Amaral, 2005). We did not identify any major hub structures within functional modules in reexposure groups. We next assessed the way in which the modular organization of the reexposure network was related to the neurobiological organization of the addiction-relevant brain structures (Koob & Volkow, 2016). Control animals exhibited spontaneous “resting-state” functional connectivity with all structures and connections sharing virtually the same state. Notably, in binge-drinking animals hardly any changes occurred, leaving the brain at functional plateau. Alcohol reexposure resulted in a major reduction in a number of links between these structures and fragmentation of functional correlations. The strongest connections among selected brain structures were found for BST and VTA that shared the same module, but also strongly correlated in terms of c-Fos activation. Of note, projections from neurons of BST to VTA are crucial to behaviors related to reward and motivation (Jalabert *et al*, 2009). Interestingly, another strong functional correlation was observed between GPe and BLA, and in particular BLAp, but not BLAa or BLAv, suggesting a dissociable role for posterior BLA in alcohol reexposure. Although much less is known of the function of the BLAp relative to BLAa, its role in cue-triggered alcohol-seeking behavior has been suggested (Millan *et al*, 2015). In humans, alcohol can change GPe activity by decreasing neuronal firing rates, suggesting that the GPe may have a central role in explaining impulsive behaviors during binge drinking (Fede *et al*, 2020). BLAp and GPe do not share any direct projection pathway, but they are indirectly connected via striatal neurons (Yager *et al*, 2015). Another strong connection was formed between HIP, SNc and CeA. The HIP is critical for the acquisition of new factual information and the formation of new memories about personally emotionally relevant experienced events (Eichenbaum, 2004), whereas CEA is related to memory consolidation for emotionally arousing events (Cahill *et al*, 1995; McGaugh *et al*, 1996). Furthermore, SNc dopamine neurons are involved in learning to predict which behavior will lead to a reward (Keiflin & Janak, 2015). Surprisingly in our study ACB was not found to be functionally connected with any of other analyzed structures.

The present study employed a novel behavioral paradigm that was based on the IntelliCage system to investigate binge-like drinking in mice. We used a high-throughput method that was based on brain clearing followed by light-sheet microscopy imaging to assess the whole-brain response to alcohol. Specific brain structures were shown to specifically respond to alcohol craving. Alterations of overall functional brain organization, including an increase in modularity and brain network disruption, in the context of alcohol craving were also observed.

## Materials and Methods

Additional details of all procedures described below are available in the Supplementary Methods and Materials.

### Behavioral training

The experiments were performed using female C57Bl6 mice that were divided into the four groups: control (CTRL group; *n* = 5), binge-like drinking (BD group; *n* = 5), cue reexposure (CR group; *n* = 5) and alcohol reexposure (AR group; *n* = 5). We used an animal model of binge-like alcohol drinking (Thiele & Navarro, 2014) that was adapted to the IntelliCage system. The procedures were performed according to the Polish Animal Protection Act, Directive 2010/63/EU, and approved by the Local Ethics Committee (permission no. 128/2016).

### Optical tissue clearing and whole-brain c-Fos imaging

Optical tissue clearing was performed using iDisco+ method (Renier *et al.*, 2016; Renier *et al.*, 2014). Brains were imaged using a light-sheet microscope (Pawlowska *et al.*, 2019; Stefaniuk *et al*, 2016).

### Computational analysis

We developed a workflow that included image registration techniques and cell detection algorithms. We took advantage of multichannel acquisitions that provided both functional and anatomical information. For each c-Fos-positive nucleus, a local maximum intensity was determined. The sum of the signal for each structure was next overlaid on its volume (in mm^3^) to create a signal density map.

### Statistical analysis

The results are expressed as mean ± SEM. Analyses were conducted using Prism 8.4.2 software (GraphPad, La Jolla, CA, USA). Differences between experimental groups were considered significant if the type 1 error was less than 5%. For comparisons of the c-Fos signal density between structures, we first performed a mixed-effect linear model following by the analysis of variance (ANOVA). The between-group differences were assessed by the two-way paired *t*-tests. Obtained *p* values were corrected for multiple comparison to *q* values (Ryabinin *et al.*, 1997). The tests were performed in Python 3.7 using the Pingouin 0.3.3 statistical package and custom-written scripts (source code and data can be found at https://github.com/pawlowska/pairwise_tests_pingouin). The network analysis was conducted using igraph package for R statistical environment; figures were prepared in Cytoscape 3.8.2 software.

## Acknowledgments

This work was supported by a TEAM grant (POIR.04.04.00-00-1ACA/16-00). MS was supported by a National Science Centre grant (2019/35/B/NZ4/04077). MP was supported by a National Science Centre grant (UMO-2015/17/B/NZ4/02631). PM and DW were supported by an ERA-NET NEURON grant from the National Centre for Research and Development (ERA-NET-NEURON/17/2017). The authors thank Prof. Kasia Radwanska and Dr. Anna Beroun for constructive criticism of the manuscript and Michael Arends for proofreading,

## Author contributions

MS and MP designed experiments and wrote the manuscript, MS and KN performed behavioral experiments, MS performed imaging, MP constructed a light-sheet set-up and performed image and computational analysis, LB and DL optimized the protocol and performed the clearing, PM, SB, NJ, ZZ and MB performed computational analysis, DW and LK supervised the work

## Data Availability Section

Datasets and codes generated and used in the reported study are listed in Supplemental Information and attached as a Supplementary File.

