## Supplemental methods for "A whole-brain imaging-based systems approach to understand origin of addiction in binge-like drinking model"

**Supplemental Information**

**Methods and Materials**

*Behavioral training*

All experiments were performed using female C57Bl6 mice (n=20) with *ad libitum* access to food, in 12:12 light to dark photoperiod. Mice were divided into four groups: water control (CTRL, n=5), binge-like drinking (BD, n=5), cue reexposure (CR, n=5) and alcohol reexposure (AR, n=5). We used an animal model of binge-like drinking of alcohol (Thiele & Navarro, 2014) adapted for IntelliCage training system. IntelliCage is a homecage equipped with fully automated corners with drink distributors. For our video showing IntelliCage, please check the movie under this [link](https://vimeo.com/486855508) (<https://vimeo.com/486855508>). The system allows for observation of mice 24/7 and the detailed analysis of their behavior in social groups. Mice were monitored individually by means of subcutaneously implanted transponders detected by a corner sensor. During each visit in a corner, a nosepoke to the door or/and lick of the bottle was noted. First, animals underwent the cage and nosepoke adaptation period (7 days). Next, they had access to 20% alcohol in drinking-in-the-dark model – alcohol was available only 2 hours per day, during the first 3 hours into the dark phase of the diurnal cycle. In the first part of the experiment (3 weeks) animals had access to alcohol in both active corners every day except for the weekends. For the next two weeks, we limited access to alcohol to one corner with concurrent access to water in the other corner. Mice were next randomly divided into groups. BD animals were not challenged with a long-term withdrawal, and their brains were collected at the end of the last drinking session. AR and CR animals had ceased access to alcohol for nine days and then were reexposed to either visual cue previously associated with alcohol (green LED light over the door, Fig. 1B) or access to alcohol matched with the cue. Control animals were kept separately and had access only to drinking water.

Amount of alcohol consumed (g/kg of body weight/day) was calculated on the basis of amount of alcohol consumed during the whole period. Average lick volume was established to 1.94 ± 0.2 μl. The following formula was used: (*number of licks per day* × *lick volume* × *alcohol concentration* × *0.79 g/ml)/animal weight*.

*Optical tissue clearing and light-sheet microscopy imaging*

To perform whole-brain imaging we first performed optical tissue clearing to render the tissue transparent [for review see (Matryba *et al*, 2019)]. Optical tissue clearing significantly reduces light scattering in the tissue, thus increasing the imaging depth by an order of magnitude or more. We used iDISCO+ technique (Renier *et al*, 2014), an organic solvent-based clearing method, where water is removed from tissue and lipids are dissolved, giving a high level of transparency, that was previously shown to enable quantitative c-Fos imaging in the whole mouse brain (Renier *et al*, 2016). In brief, animals were first anesthetized with isoflurane and then sacrificed with an injection of a lethal dose of sodium pentobarbital. Mice were perfused using 0.01 M PBS (pH 7.4)/heparin (5 IU/ml final concentration) and 4% PFA/0.01M PBS. Brains were isolated and postfixed in 4% PFA for 12 hours. Next, samples were dehydrated with an increasing concentration of methanol, bleached in H_2_0_2_ and again rehydrated. Next, samples were immunostained for c-Fos. We followed the most recent protocol provided by [https://idisco.info/](https://idisco.info/%20) (version from December 2016). Briefly, samples were incubated in Permeabilization Solution in 37°C for 2 days, blocked in Blocking Solution in 37°C for 2 days; incubated with primary antibody (anti-c-Fos 1:2000, Synaptic Systems, #226-003) in Blocking Solution/5%DMSO/3% Donkey Serum in 37° for 2 days, washed in PTwH (PBS/2%Tween-20/1% heparin) until the next day. Next, samples were incubated with secondary antibody (anti-rabbit-AlexaFluor 647, Molecular Probes, #A27040) in PTwH/3% Donkey Serum in 37° for 2 days. After incubation, samples were washed in PTwH five times and left until the next day at room temperature. All steps were performed on a rotary tube suspension mixer. Then samples were dehydrated again with methanol and dichloromethane as the last step. Finally, brains were cleared using dibenzyl ether. Such prepared samples were next imaged in the light-sheet microscope (LSFM) at least 24 h after clearing which was sufficient time for the stabilization of fluorescence (Pawlowska *et al*, 2019; Stefaniuk *et al*, 2016). We performed LSFM imaging on our homebuilt setup that was already used for imaging of cleared samples including the brain (Matryba *et al*, 2018; Stefaniuk *et al.*, 2016). For each brain, we acquired two channels – c-Fos (638 nm excitation) and background autofluorescence (488 nm excitation) that was later used to annotate the brain with the atlas. The microscope was equipped with a 4×, 0.3 NA immersion objective corrected for refractive indices up to 1.6 (LaVision Biotec LVMI-Fluor 4×/0.3) and a sCMOS camera with 2048 × 2048 pixels (Hamamatsu Orca 4) which corresponds to a pixel size of 1.45 × 1.45 µm. In this setup, the light sheet is generated statically with a cylindrical lens which focus is relayed into the back focal plane of the illumination objective (Gualda *et al*, 2013; Pitrone *et al*, 2013). A slit placed before the cylindrical lens enables adjusting the light-sheet thickness. The thickness was set to about 10 µm as a compromise between the axial resolution and illumination uniformity along the propagation direction of the light. To further improve the resolution of the c-Fos data, we limited the field of view used for imaging and used a z-step of 4 µm.

*Data Preprocessing*

The output of the light-sheet microscope is a series of single-channel TIFF stacks: one per channel and per tile at the native resolution of the camera, 1.45 μm/pixel, with a few hundred slices separated by 4 μm for the c-Fos channel and 10 μm for the autofluorescence channel, respectively. Because the autofluorescence data was taken with a larger field-of-view, an additional preprocessing step was required to remove vignetting artifacts resulting from a finite height of the light sheet. This was done by adapting the BaSiC plugin (Peng *et al*, 2017) to large 3D data. First, a subset of data was taken by taking every 20^th^ slice and based on this subset, the flat-field and dark-field corrections were calculated. Next, the entire dataset was corrected slice by slice. The data was then stitched using Big Stitcher Image-J plugin (Horl *et al*, 2019). We also tested Terastitcher (Bria & Iannello, 2012). However, while both plugins worked well for good quality data, Terastitcher offers less flexibility in situations when there is non-uniform tile spacing or a missing tile caused by hardware problems during acquisition. Finally, we removed absorption and scattering artifacts with VSNR algorithm (Escande *et al*, 2017).

*Computational Analysis*

*Infrastructure*

To facilitate the image processing, registration, and analysis, we first converted the series of TIFF files into a multiresolution pyramid file format based on the HDF5 container using the LSFMPy toolkit (<https://pypi.org/project/lsfmpy/>). Subsequently, the images of both channels (i.e., autofluorescence and c-Fos signal intensity) were exported at 25 μm isotropic resolution. Additionally, to mitigate the chance of gross misregistration, the CCF3 template was masked, for each experimental brain separately, to exclude these parts of each given hemisphere that were not imaged, or were damaged during the experimental handling (e.g., olfactory bulb or brainstem).

*Tissue segmentation*

To facilitate the registration procedure we trained a 2-dimensional Deep Convolutional Neural Network (DCNN) based on the UNet architecture (Ronneberger, 2015) to perform automated segmentation of several anatomical brain structures on the autofluorescence channel image. Training datasets for the DCNN were created by manual annotation of subsections of the images in the three cardinal planes (coronal, sagittal, and horizontal) using ITKSnap v. 3.8.0 software <http://www.itksnap.org>, (Yushkevich *et al*, 2006). The trained network allowed us to automatically identify the outline of the entire brain as well as the main white matter tracts, the dentate gyrus, and characteristic parts of the cerebellum: white matter, granular cells layer, and molecular cells layer. Note that the selection of these areas was dictated solely by the performance of the subsequent registration procedure and is unrelated to the scientific goals of the study.

*Registration*

The registration process was carried out with Advanced Normalization Tools (ANTs 1.9.4, (Avants *et al*, 2011), RRID:nlx 75959). First, the two LSFM image channels (autofluorescence and c-Fos) were affinely aligned based on their outlines, with Mattes Mutual Information (MI) as a similarity metric. Then the autofluorescence channel and CCF3 template were aligned by, first, the center of mass, followed by an affine alignment. Finally, the autofluorescence channel image and the template were non-linearly registered with Symmetric Diffeomorphic Normalization (SyN, (Avants *et al*, 2008) algorithm (gradient step of 0.25, regularization by Gaussian smoothing with σ = 1.75 voxels for similarity gradient, and σ = 0.75 voxels for the displacement field, maximum number of iterations: 200×200×100). Three, equally weighted, pairs of images were used to simultaneously drive the registration. (1) For the intensity images (i.e. the autofluorescence channel and the CCF3 template) we used the normalized correlation coefficient (Avants & Gee, 2004) with the window size of 5 voxels. (2) The overlap between the segmented brain areas in an experimental brain and the CCF3 template was assured by the use of Point-Set Estimation (PSE) (Avants *et al.*, 2011) metric with an exhaustive (100%) sampling of the labelled voxels. Finally, (3) the mean squared difference (MSQ) between the outlines of the brains being coregistered was used to further increase the registration reliability. Upon warping the autofluorescence and c-Fos intensity images to the CCF3 template, the results were inspected in ITKSnap to visually confirm the quality of registration.

*c-Fos-positve nuclei detection*

To identify the locations of individual c-Fos-positive nuclei we used a 3D UNet DCNN. The network was trained on a procedurally generated batch of images resembling the actual images of the c-Fos signal channel. For instance, the simulated stained nuclei were represented as 3D Gaussian blobs and displayed variability in sizes, intensities, and densities which corresponded to that of the experimental data. We also modelled typical artifacts of the LSFM imaging process such as deformation of the objects along the z-plane of the microscope setup due to the hyperbolic profile of the light sheet along the propagation direction, resulting from diffraction. The network was trained to convert the input c-Fos signal image to a nuclei density map. The prediction was performed on a series of subvolumes of the c-Fos channel image exported from the LSFMPy container. Between 30 and 40 thousands of subvolumes of the physical size of 185.6×242.0×371.2 µm represented at an isotropic resolution of 1.45 μm per voxel were necessary to cover the extent of an entire brain hemisphere. The c-Fos-positive nuclei were identified by local maxima detection on the resulting density maps, and their physical coordinates in the c-Fos channel as well as their maximum intensities were recorded. Heat maps were generated for each case by marking the locations of each detected nuclei on a 25 μm isotropic canvas in the space of the c-Fos signal channel. The contribution of each nucleus to the heat map was equal to its maximum intensity. To correct for periodic artifacts arising due to lower sensitivity of the detection at the edges of input image patches, the heat maps were smoothed with a Gaussian kernel (σ = 35 µm) followed by a Fourier filtering. The maps were then registered to the space of the CCF3 template using the previously computed spatial transformations. Finally, based on template segmentation mapped to the c-Fos channel, signal density was determined for each brain structure defined in the reference atlas (sum of the signal within a given structure divided by its volume).

For statistical analysis signal densities for brain structures were aggregated at depth up to level 6 (Figure 3, Supplementary Figures 2-6, Supplementary Dataset 1) and up to level 5 in Figure 4 and Supplementary Figures 7-10. After excluding missing values, this left 169 (level 5) and 258 (level 6) structures that were considered in the analysis.

*Correlation maps and modularity analysis*

The data consisting of signal densities for each brain structure and each mouse was further analyzed with scripts written by us in R. Source code and data can be found at <https://github.com/pawlowska/Stefaniuk_et_al_correlation_analysis>. Since the version of the atlas, we used is very detailed, containing in total 1325 structures divided into layers (see Supplementary Table 1, tab “depths_struct_all_ABA”) and not all structures were identified in terms of c-Fos signal detected (see Supplementary Table 1, tab “depths_struct_det_ABA”), we aggregated the data for clarity to show structures at depth up to 5. After excluding missing values, this left 169 structures that were considered in the analysis. Whole-brain clearing and imaging is a complex procedure and with 20 animals, it is difficult to avoid differences between specimens due to clearing quality or fluorescence quenching by the clearing reagents. These factors increase the standard deviation of the measured signal within a group. However, if a specimen has suffered from fluorescence quenching, this will result in a lower signal density in all brain structures simultaneously, so these global effects can be circumvented by considering correlations between pairs of structures instead of comparing means and their errors. For this, we calculated Pearson correlations between pairs of structures within each group. First, we wanted to look at the correlation within the broader anatomical regions of the brain. We divided the structures into categories that correspond to the following subtrees of the atlas tree: cortical plate (CTXpl), cortical subplate (CTXsp), striatum (STR), pallidum (PAL), thalamus (TH), hypothalamus (HY), midbrain (MB) and hindbrain (HB), see Supplementary Table 1, tabs “AtlasTree”. The cerebellum was left out in the analysis. Next, we looked at how the structures group together according to their signal density. A high correlation between two structures can be accidental; for two structures to be considered ‘similar’, we want their respective correlations with every other structure in the brain to be similar. This can be described by complete Euclidean distances between rows of the correlation matrix (computed using *dist* function from the *stats* package in R).  Based on these distances, the brain structures were then grouped using the complete hierarchical clustering algorithm (*hclust* in R). Hierarchical clustering works by iteratively grouping together structures and then clusters of structures with the smallest distance, corresponding to the largest similarity.  The result can be presented as a dendrogram in which the height at which any two structures are first connected corresponds to how similar they are. To compare the dendrograms obtained for different experimental groups, we plotted the number of clusters depending on relative tree height. The hierarchical cluster dendrograms were trimmed at half the height of each given tree to split the dendrogram into specific modules (see Supplementary Table 2).

*Network measures of brain connectivity*

The understanding of a complex system such as brain connectivity requires a multidisciplinary approach known as network analysis. A network represents real-world connectivity and is defined by a collection of nodes (vertices) and links (edges) between pairs of nodes (Rubinov & Sporns, 2010). In brain networks presented in this study nodes represent brain regions, while links represent functional connections, based on the modules identified on hierarchical cluster dendrograms derived from complete Euclidean distances analysis. The measures of centrality were used to identify potentially important brain regions and their interaction between and within functional modules, and further heuristic classification of nodes into distinct functional groups (Guimera & Nunes Amaral, 2005). In the functional network analysis, a positive correlation (Chen *et al*, 2011; Giove *et al*, 2009; Meunier *et al*, 2009; Murphy *et al*, 2009) thresholded to a value higher than 0.75 of the Pearson correlation coefficients (Orsini *et al*, 2018; Wheeler *et al*, 2013) were used. The centrality was assessed using the measurement of within-module degree z-score (WMD) and the participation coefficient (PC). The role of each node was defined by the values of both WMD and PC of that module (Guimera & Nunes Amaral, 2005).

The within-module degree z-score is a localized degree centrality, i.e. the importance of module for within-module connectivity, which provides a measure of the extent to which node is connected to other nodes in the same functional module. WMD is calculated as:

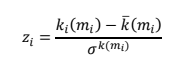

where *k* is a within-module degree, *mi* is the module containing node *i*, *ki(mi)* is the within-module degree of *i* (the number of links between *i* and all other nodes in *mi*), and *k(mi)* and *k(mi)* are the respective mean and standard deviation of the within-module mi degree distribution.

The participation coefficient provides a measure of how the connections of a node are distributed. PC value close to 1 indicates that connections are uniformly distributed among all the functional module, while PC value close to 0 indicates that connections are within its module. The PC is calculated as:

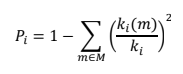

where *M* is the set of modules, *ki(m)* is the number of links between *i* and all nodes in module *m*, and let *ki* is a total degree of the node.

Nodes can be classified by their role based on the measure of its within-module degree and participation coefficient (Guimera & Nunes Amaral, 2005). All nodes with WMD equal or higher than 2.5 are considered as hubs, while nodes with WMD lower than 2.5 as non-hubs. Further classification is based on values of PC:

- peripherals – with low WMD and PC values
- connectors – with low WMD but high PC values, they connect modules having a low number of connections within their module
- provincial hubs – with high WMD and low PC values, are likely to play an important part in the facilitation of modular segregation
- connector hubs – with high values of both WMD and PC, are likely to facilitate global intermodular integration

*Statistical analysis*

All results are expressed as mean ± SEM. Behavioral analyses were conducted using GraphPad Prism, version 8.4.2 (GraphPad Software, Inc., La Jolla, CA). Differences between the experimental groups were considered significant if the type 1 error was less than 5%. To compare signal densities of c-Fos signals, we compared subjects using mixed ANOVA with Group as a between-subject factor and anatomical regions (structures) as a within-subject factor. Next we performed two-way pairwise t-Tests to obtain *p* values of between-group differences. P-values were corrected for multiple comparison to *q* values to control for false-discovery rate (Benjamini & Hochberg, 1995), taking into account comparisons between all the possible group pairs (6) and 169 structures, in total 1014 comparisons. Tests were performed in Python 3.7, using Pingouin (Vallat, 2018) version 0.3.3 statistical package and custom-written scripts. Source code and data can be found at <https://github.com/pawlowska/pairwise_tests_pingouin>. The network analysis was conducted using igraph package for R statistical environment; figures were prepared in Cytoscape 3.8.2 software.

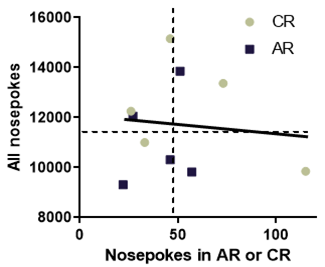

**Figure S1**

No correlation between overall mouse activity (as defined by cumulative number of nosepokes) and nosepokes in reexposure to cue (CR) or alcohol (AR). Individual squares or circles refer to individual mice. R^2^=0.0078, p=0.8073.

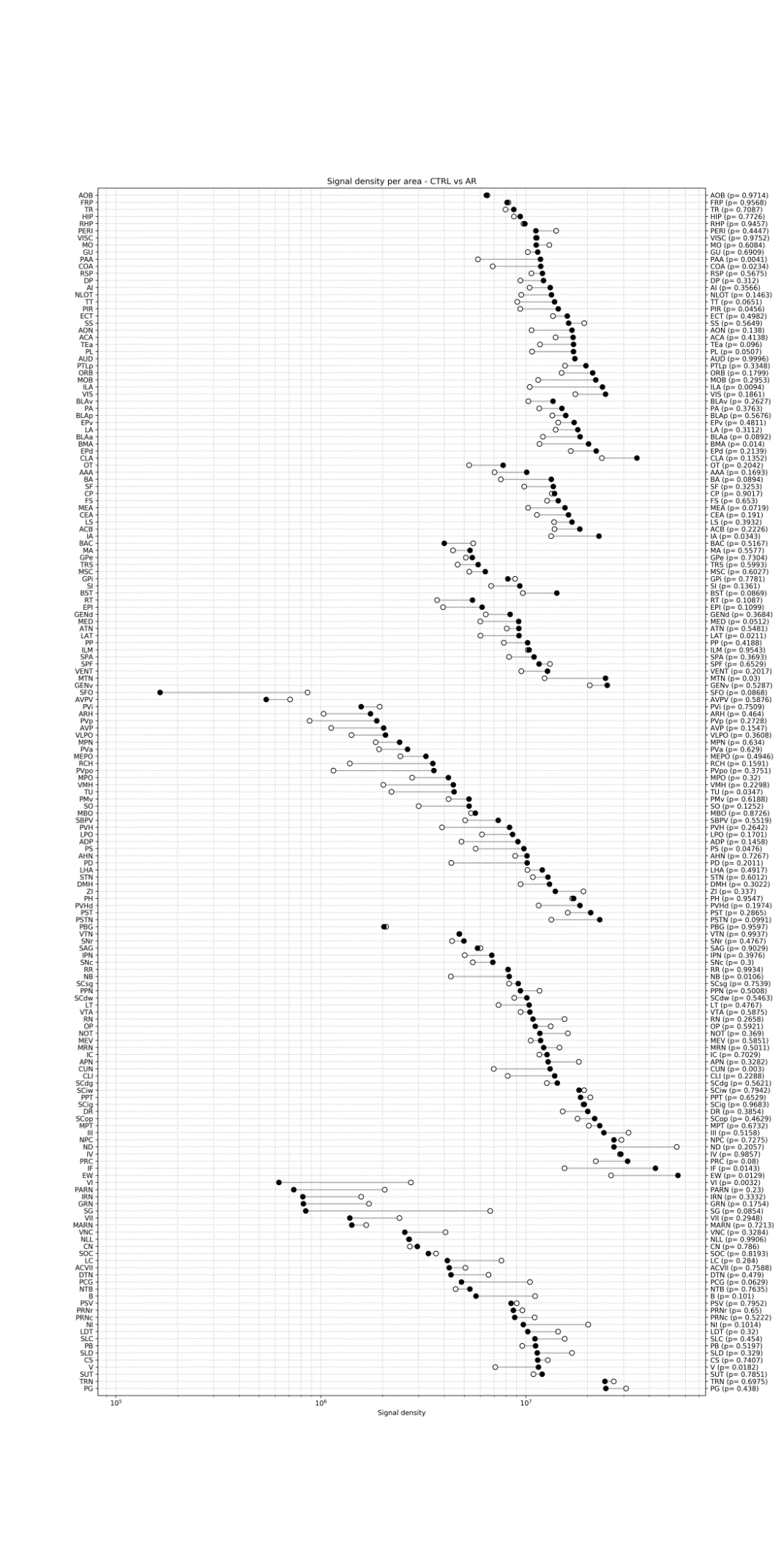

**Figure S2**

Overview of the c-Fos signal density profile in the whole brain, showing a comparison between the CTRL group and AR group. The signal density for each structure was obtained by dividing the sum of signals of individual c-Fos-positive cells in a given structure by its volume (in mm^3^). The structures are grouped in anatomical categories according to the Allen Brain Atlas. Within-category structures are shown according to decreasing signal density. The data were analyzed using one-way paired *t*-test, and resulting *p* values are in brackets next to the structure acronym. Empty circle – control animals, black circle – alcohol reexposure animals

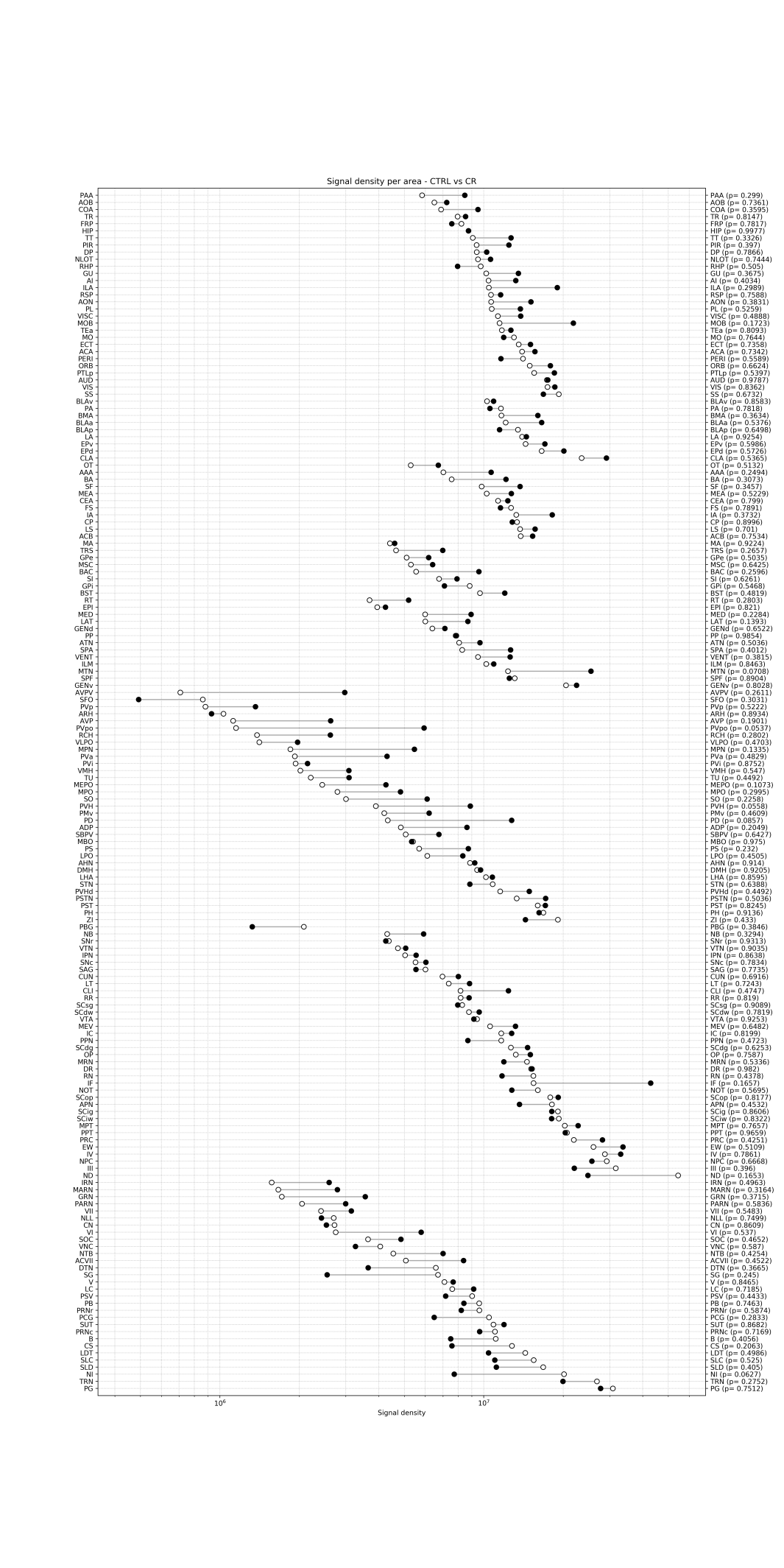

**Figure S3**

Overview of the c-Fos signal density profile in the whole brain, showing a comparison between the CTRL group and CR group. The signal density for each structure was obtained by dividing the sum of signals of individual c-Fos-positive cells in a given structure by its volume (in mm^3^). The structures are grouped in anatomical categories according to the Allen Brain Atlas. Within-category structures are shown according to decreasing signal density. The data were analyzed using one-way paired *t*-test, and resulting *p* values are in brackets next to the structure acronym. Empty circle – control animals, black circle – cue reexposure animals

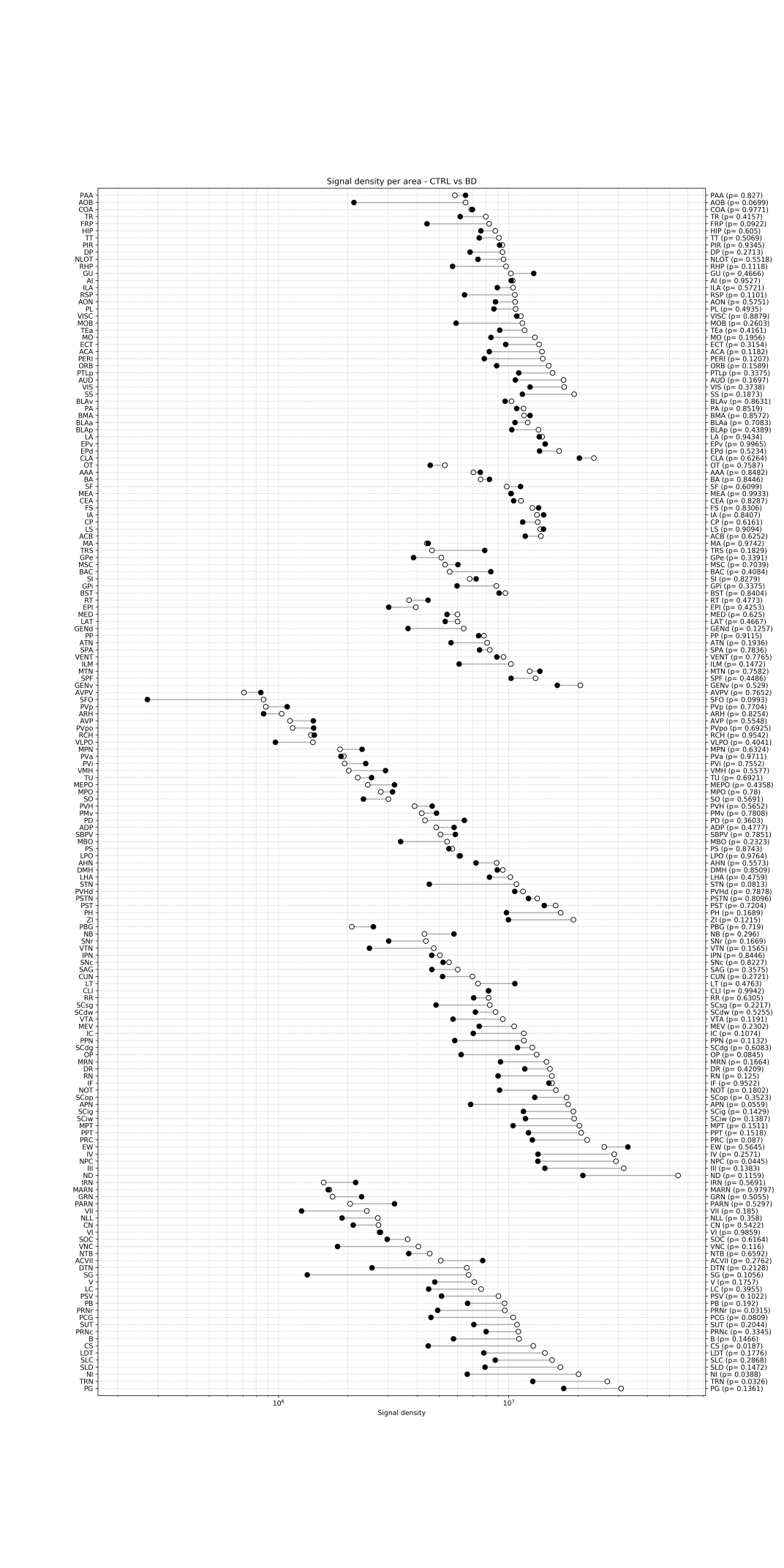

**Figure S4**

Overview of the c-Fos signal density profile in the whole brain, showing a comparison between the CTRL group and BD group. The signal density for each structure was obtained by dividing the sum of signals of individual c-Fos-positive cells in a given structure by its volume (in mm^3^). The structures are grouped in anatomical categories according to the Allen Brain Atlas. Within-category structures are shown according to decreasing signal density. The data were analyzed using one-way paired *t*-test, and resulting *p* values are in brackets next to the structure acronym. Empty circle – control animals, black circle – binge-drinking animals

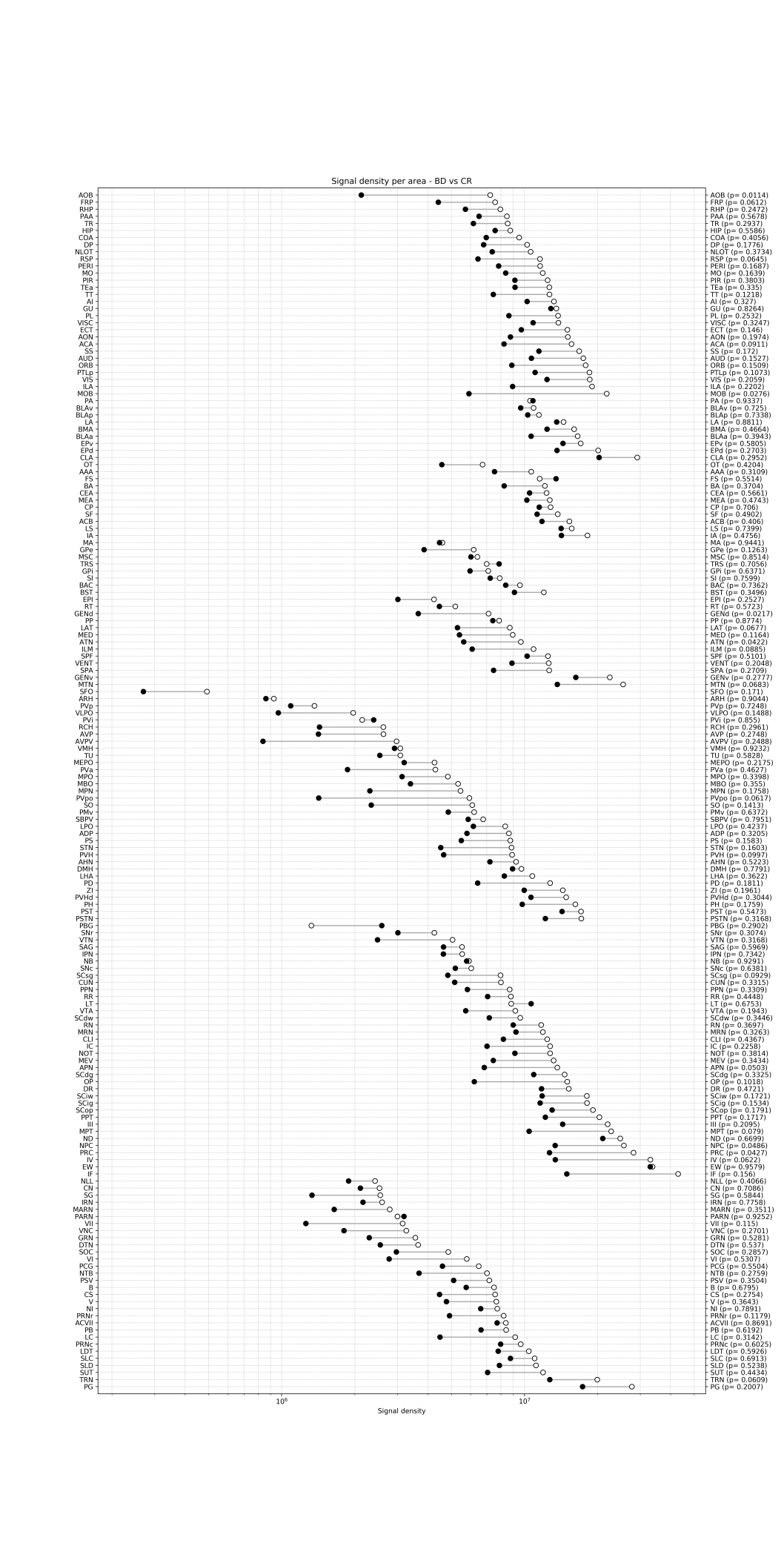

**Figure S5**

Overview of the c-Fos signal density profile in the whole brain, showing a comparison between the BD group and CR group. The signal density for each structure was obtained by dividing the sum of signals of individual c-Fos-positive cells in a given structure by its volume (in mm^3^). The structures are grouped in anatomical categories according to the Allen Brain Atlas. Within-category structures are shown according to decreasing signal density. The data were analyzed using one-way paired *t*-test, and resulting *p* values are in brackets next to the structure acronym. Empty circle – cue reexposed animals, black circle – binge-like drinking animals

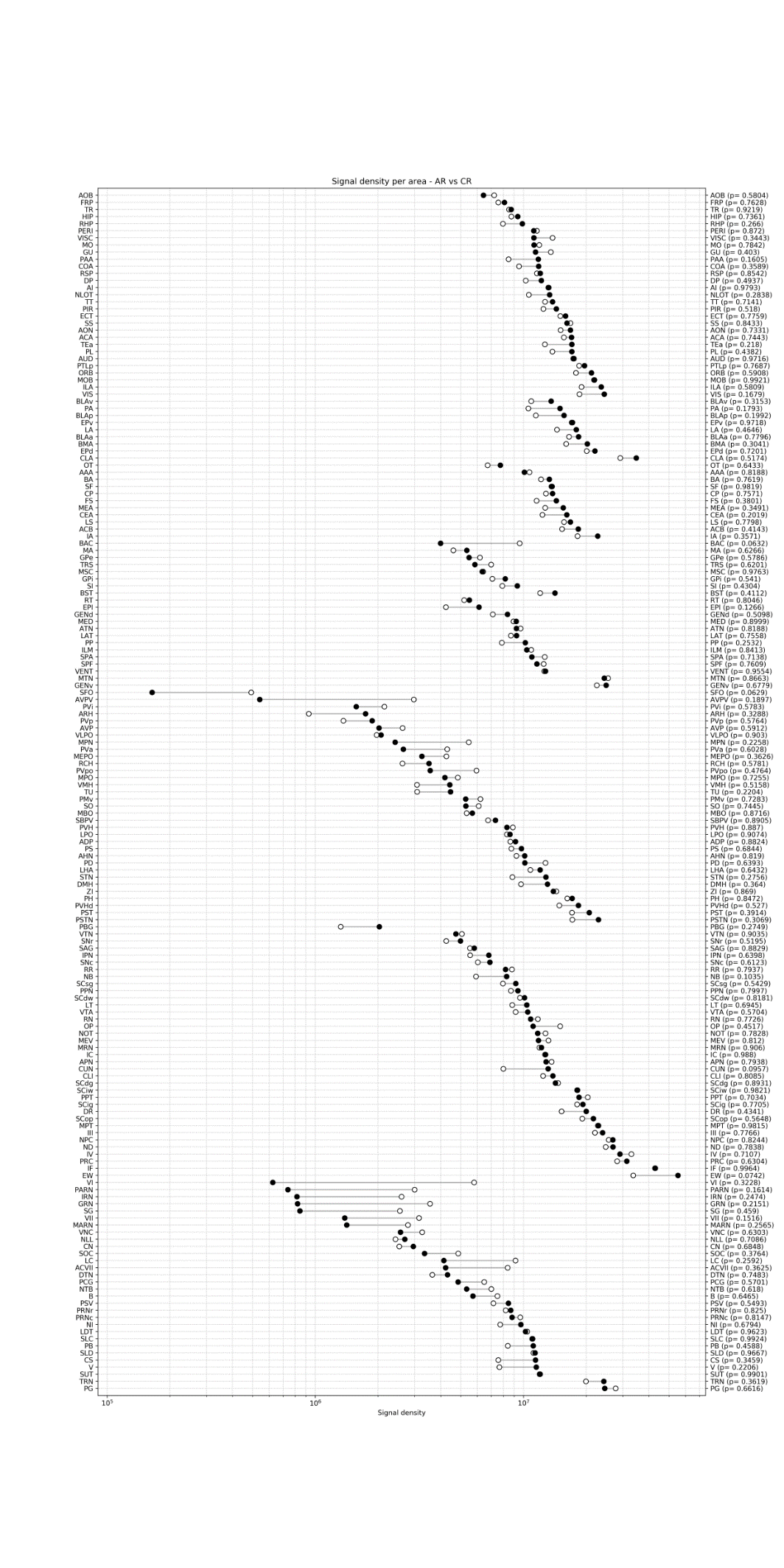

**Figure S6**

Overview of the c-Fos signal density profile in the whole brain, showing a comparison between the CR group and AR group. The signal density for each structure was obtained by dividing the sum of signals of individual c-Fos-positive cells in a given structure by its volume (in mm^3^). The structures are grouped in anatomical categories according to the Allen Brain Atlas. Within-category structures are shown according to decreasing signal density. The data were analyzed using one-way paired *t*-test, and resulting *p* values are in brackets next to the structure acronym. Empty circle – cue reexposed animals, black circle – alcohol reexposed animals

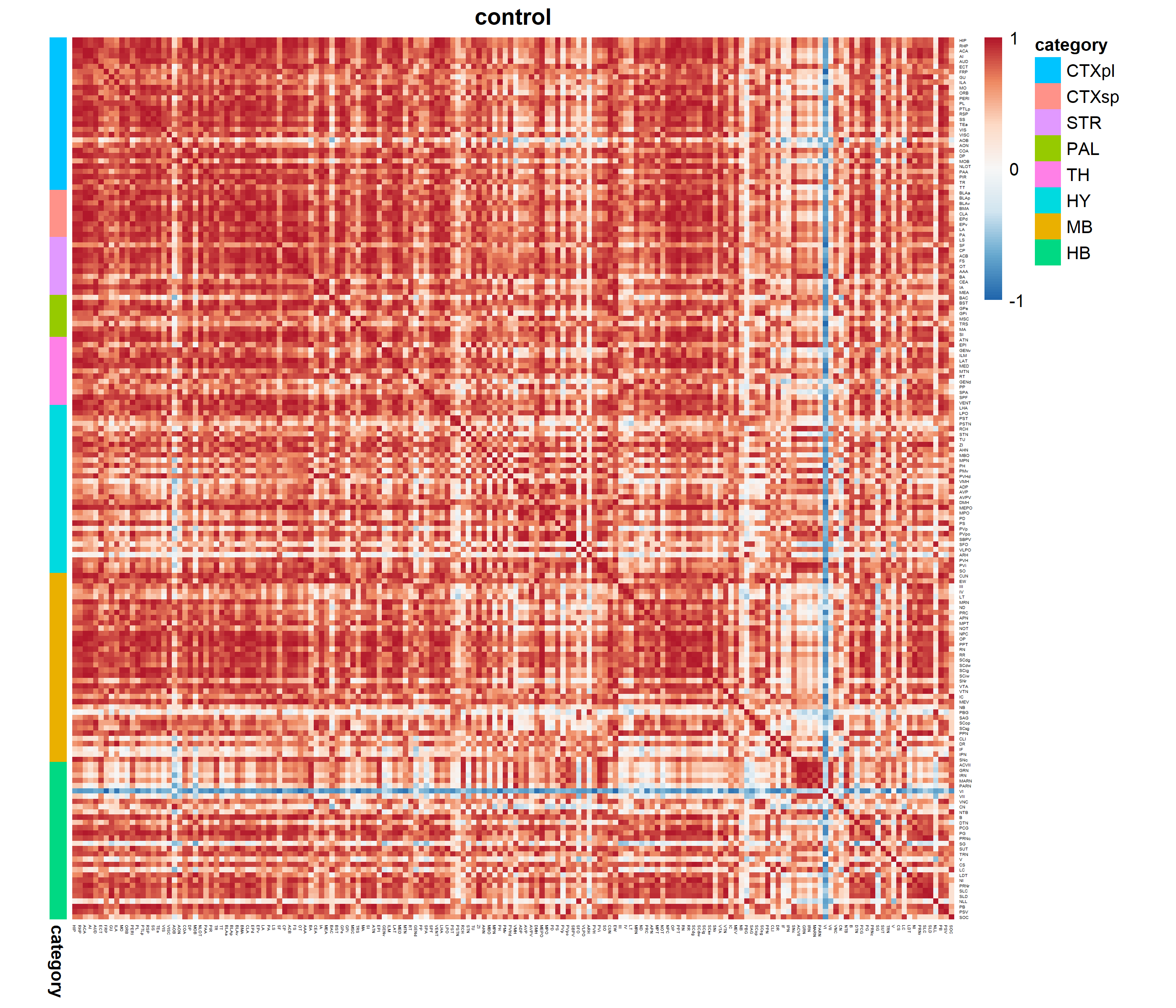

**Figure S7**

c-Fos signal density correlation map for naive control (CTRL) group having access to water only and no previous exposure to alcohol. Color coded anatomical annotations are as depicted above the heat maps. R indicates correlation strength between single structures, i.e. the warmer the color the more similar in signal density are two structures in all mice (n=5) from a given group. The colder the color the correlation is lower. CTXpl – cortical plate, CTXsp – cortical subplate, STR – striatum, PAL – pallidum, TH – Thalamus, HY – Hypothalamus, MB – Midbrain, HB - Hindbrain

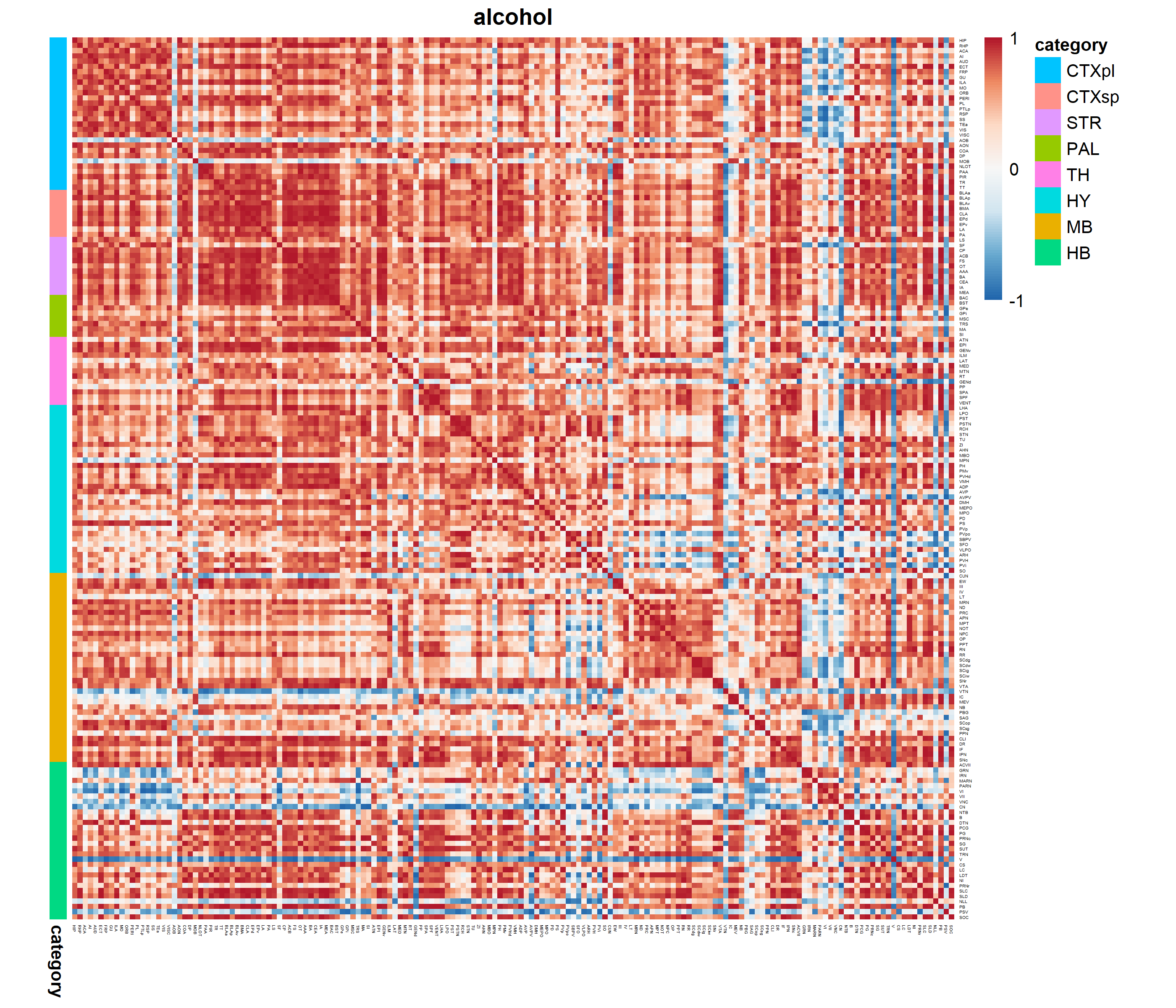

**Figure S8**

c-Fos signal density correlation map for binge drinking (BD) mice snap frozen after the last drinking session. Color coded anatomical annotations are as depicted above the heat maps. R indicates correlation strength between single structures, i.e. the warmer the color the more similar in signal density are two structures in all mice (n=5) from a given group. The colder the color the correlation is lower. CTXpl – cortical plate, CTXsp – cortical subplate, STR – striatum, PAL – pallidum, TH – Thalamus, HY – Hypothalamus, MB – Midbrain, HB - Hindbrain

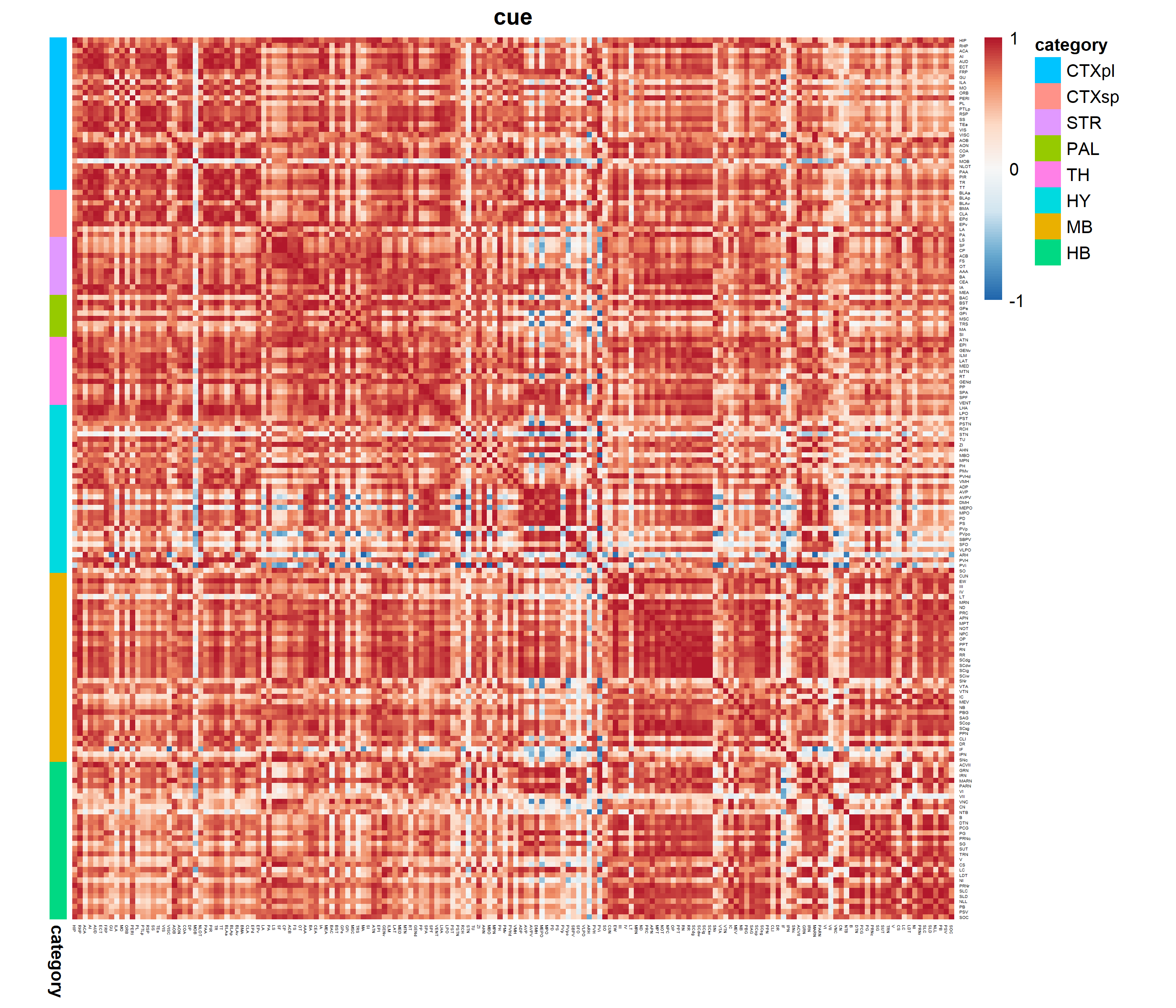

**Figure S9**

c-Fos signal density correlation map for cue reexposed (CR) animals following alcohol withdrawal. Color coded anatomical annotations are as depicted above the heat maps. R indicates correlation strength between single structures, i.e. the warmer the color the more similar in signal density are two structures in all mice (n=5) from a given group. The colder the color the correlation is lower. CTXpl – cortical plate, CTXsp – cortical subplate, STR – striatum, PAL – pallidum, TH – Thalamus, HY – Hypothalamus, MB – Midbrain, HB - Hindbrain

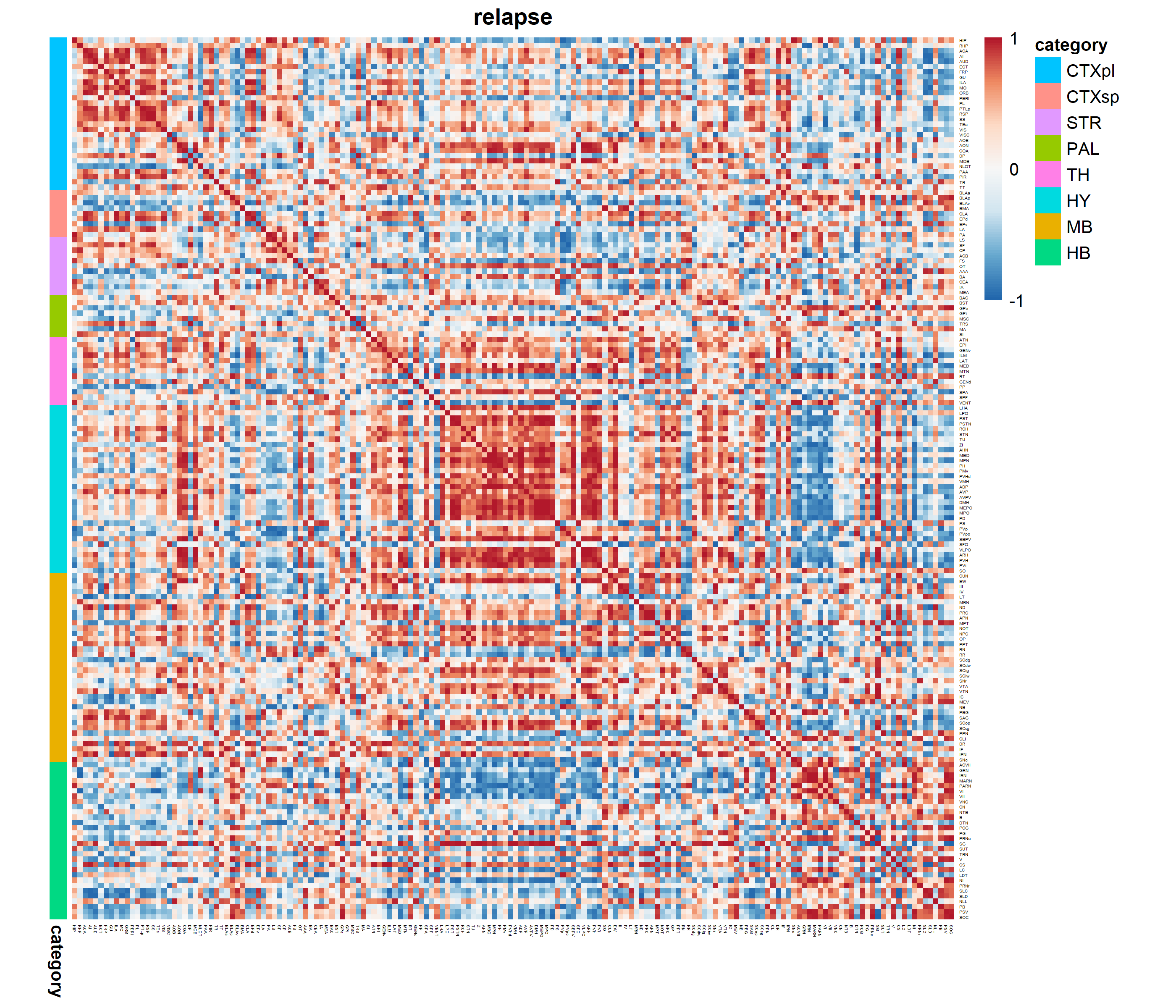

**Figure S10**

c-Fos signal density correlation map of alcohol reexposed animals following alcohol withdrawal. Color coded anatomical annotations are as depicted above the heat maps. R indicates correlation strength between single structures, i.e. the warmer the color the more similar in signal density are two structures in all mice (n=5) from a given group. The colder the color the correlation is lower. CTXpl – cortical plate, CTXsp – cortical subplate, STR – striatum, PAL – pallidum, TH – Thalamus, HY – Hypothalamus, MB – Midbrain, HB - Hindbrain

**Figure S11**

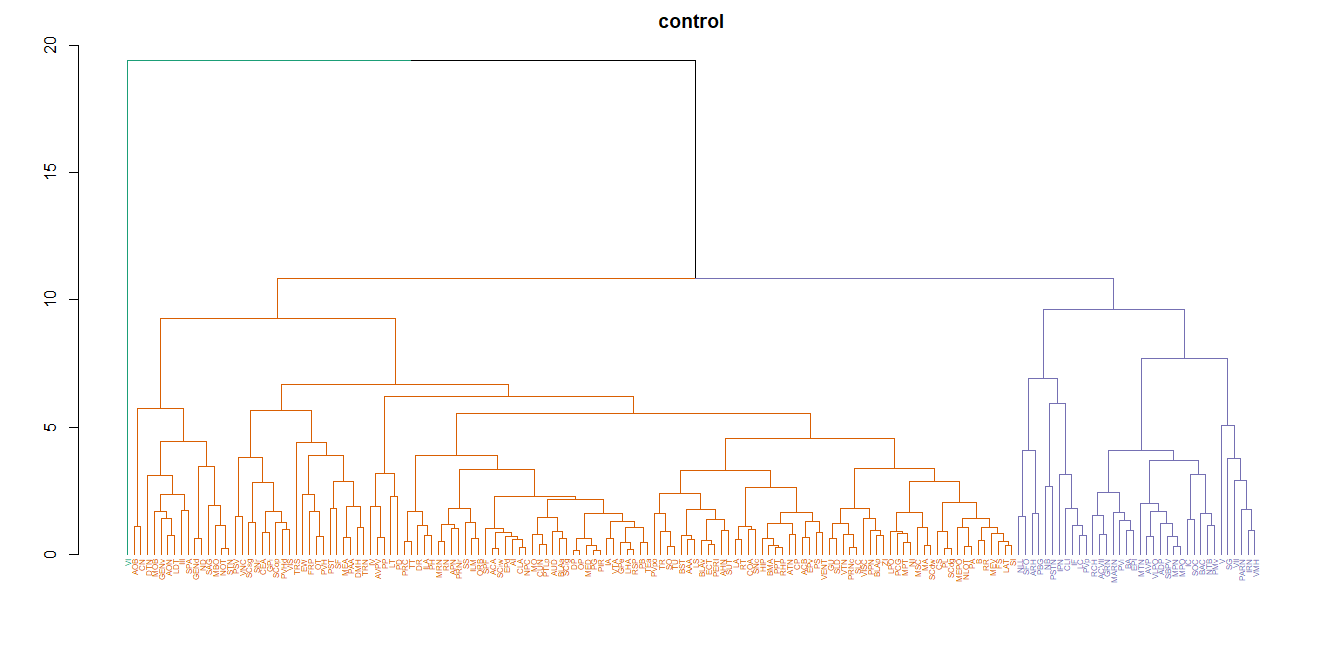

Graphical representation of structures in modules in CTRL group. Structures in the same color belong to the same module. The closer the modules within the tree, the more similar pattern of c-Fos signal density comparing to the neighbor module they share. Related to Figure 5A and Supplemental Table 2.

**Figure S12**

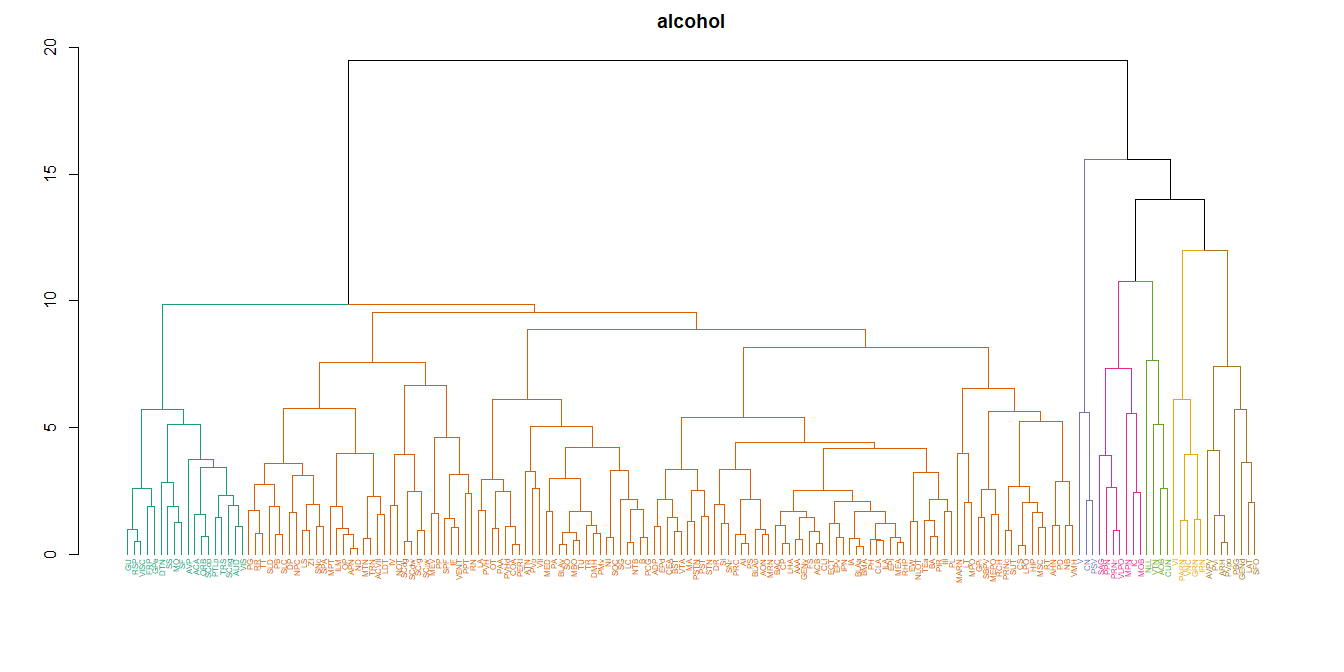

Graphical representation of structures in modules in BD group. Structures in the same color belong to the same module. The closer the modules within the tree, the more similar pattern of c-Fos signal density comparing to the neighbor module they share. Related to Figure 5B and Supplemental Table 2.

**Figure S13**

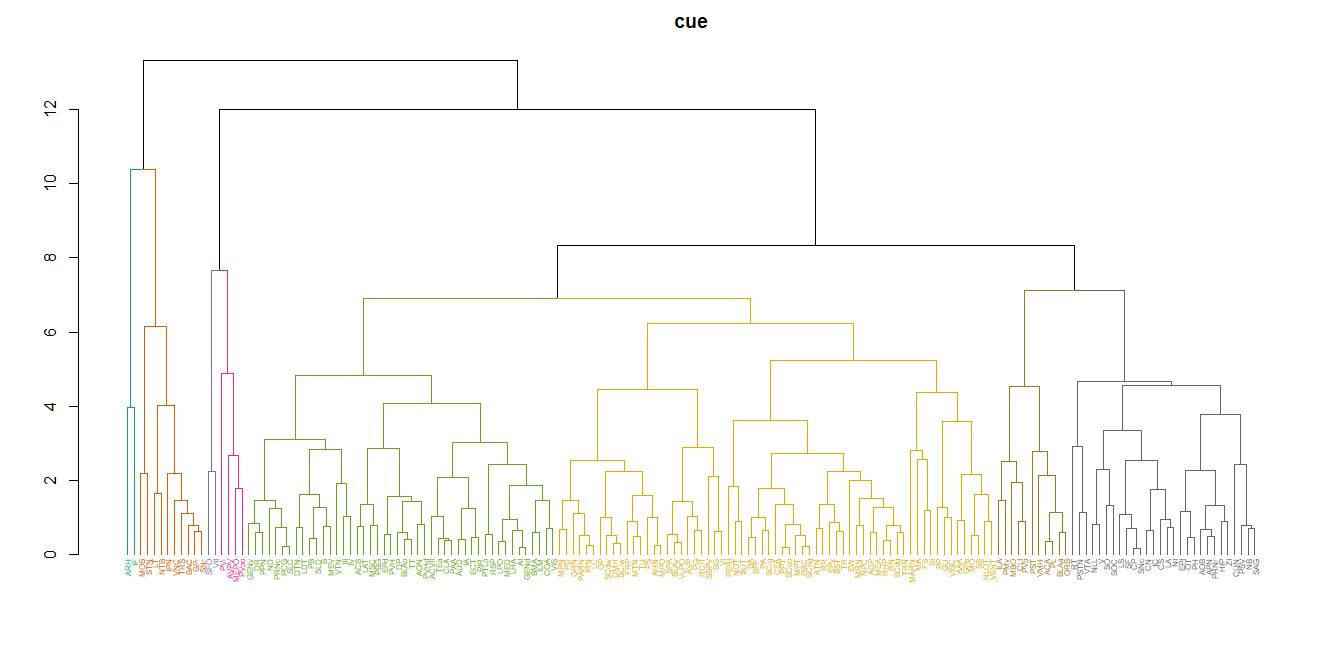

Graphical representation of structures in modules in CR group. Structures in the same color belong to the same module. The closer the modules within the tree, the more similar pattern of c-Fos signal density comparing to the neighbor module they share. Related to Figure 5C and Supplemental Table 2.

**Figure S14**

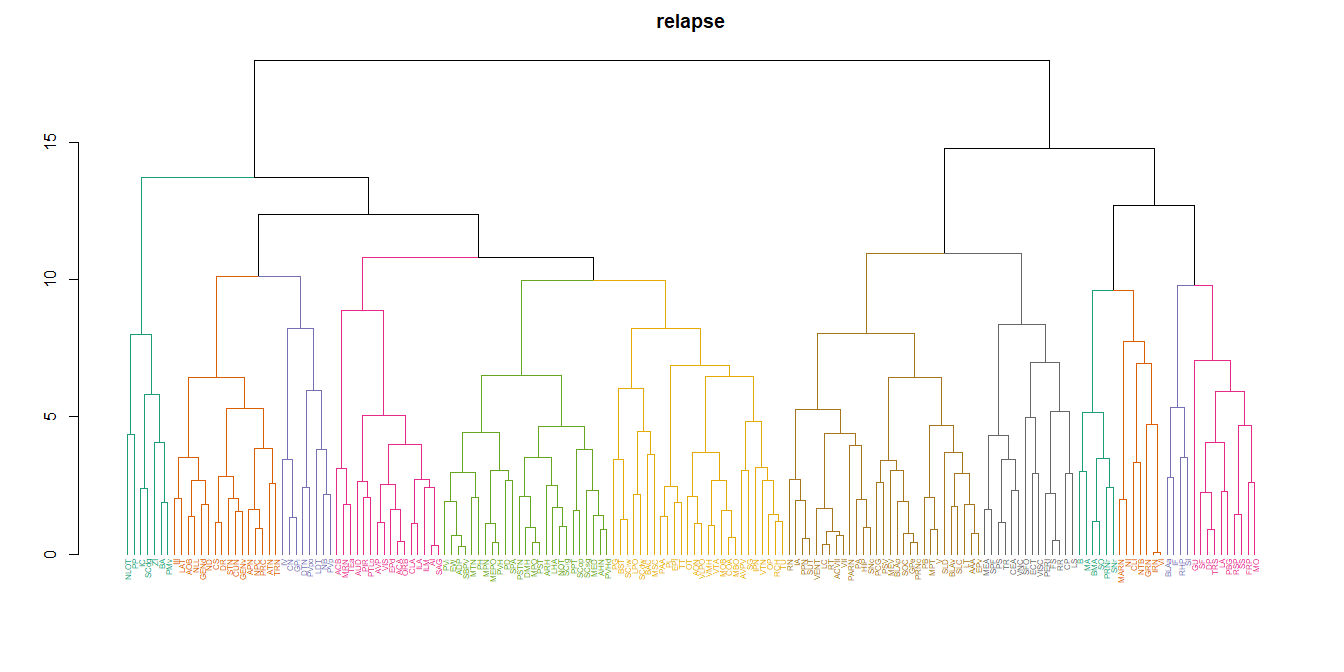

Graphical representation of structures in modules in AR group. Structures in the same color belong to the same module. The closer the modules within the tree, the more similar pattern of c-Fos signal density comparing to the neighbor module they share. Related to Figure 5D and Supplemental Dataset 2.

**Figure S15**

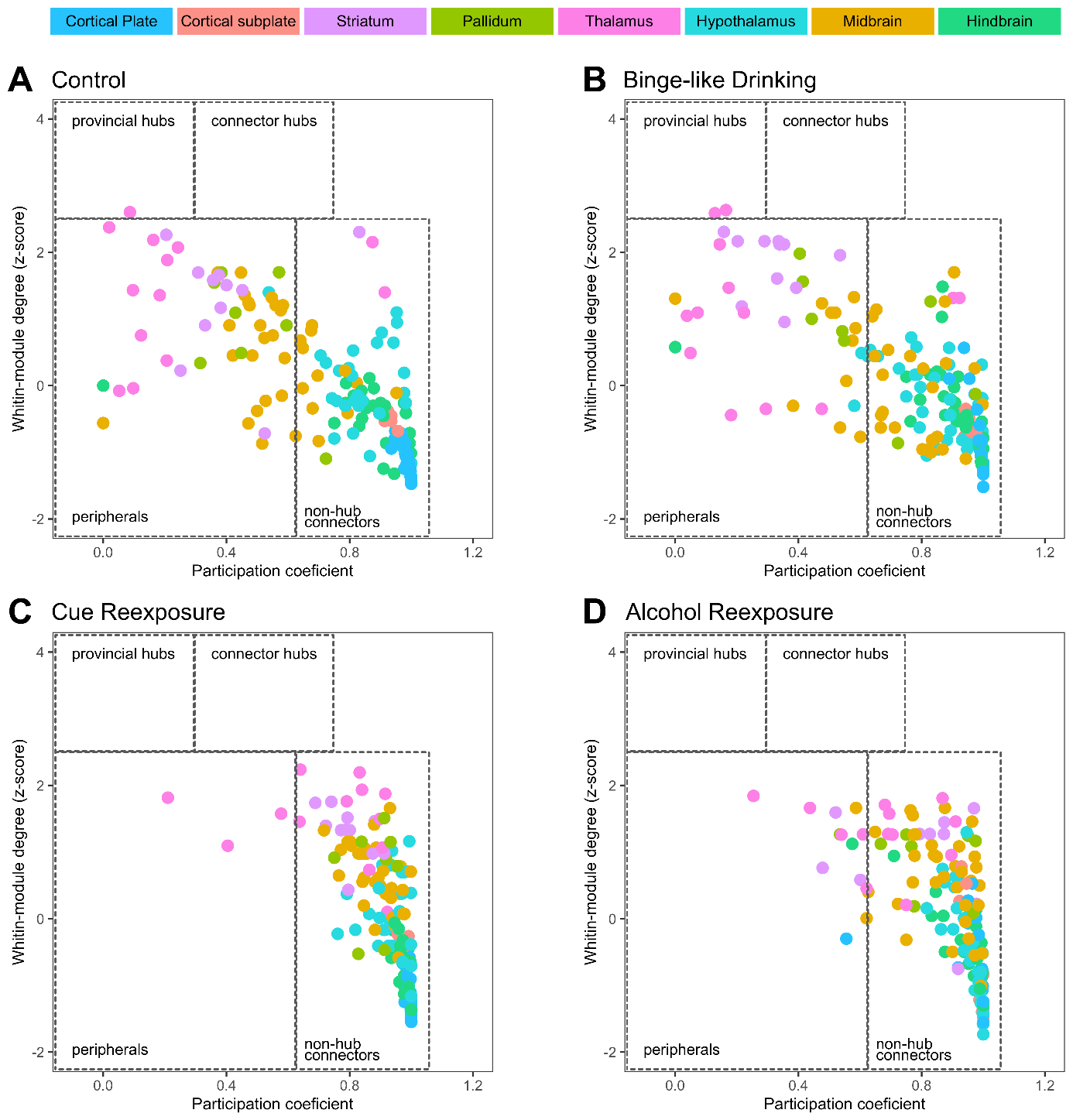

Network analysis functional cartography map showing the distribution of nodes (brain structures) in terms of participation coefficient (PC; horizontal axis) and within-module degree z-score (WMD, vertical axis). Localization in WMD–PC space classifies the node as one of the roles it plays in the network (provincial hub, connector hub, non-hub connector or peripheral node) (Guimera & Nunes Amaral, 2005). Colors indicate anatomical brain regions in which structures are located, in animals from the (A) control group, (B) binge-like drinking group, (C) cue reexposure group, and (D) alcohol reexposure group.

**Supplementary Movie 1**

Reconstruction of c-Fos positive nuclei in the hemisphere of a mouse reexposed to alcohol after withdrawal. Coronal sections represent anatomical annotations on the basis of Allen Brain Atlas. The background autofluorescence was generated by Serial 2-photon Tomography (<https://idisco.info/clearmap-2/>). Movie created in Imaris.

**Supplementary Dataset 1**

“Pairwise_2sided_depth6” tab - region statistics output of signal density data. Each region is sorted by alphabetical order. *p* values are given for each pair of groups. Mean, standard deviation, *p* values, FDR correction of the *p* values and individual sample statistics are presented for each region. Comparison was made up to anatomical level 6 of Allen Brain Atlas.

“abbrev_names_depth6” – names of structures and corresponding abbreviations made up to level 6 of Allen Brain Atlas

“depths_struct_det_ABA” – structures detected in experiment with corresponding anatomical levels (L0-L9)

“depths_struct_all_ABA” – all structures of Allen Brain Atlas sorted by anatomical levels (L0-L9)

“AtlasTree_depth5” – division into gross categories for structures up to level 5 on the basis of Allen Brain Atlas. Related to Figure 4 and Figure S7-10

“AtlasTree_depth6” tab - division into gross categories for structures up to level 6 on the basis of Allen Brain Atlas. Related to Figure 3 and Figure S2-6

“DID_depth5_signal_dens” tab – raw data of signal densities for structures up to level 5

“DID_depth6_signal_dens” tab – raw data of signal densities for structures up to level 6

**Supplementary Dataset 2**

Division to modules for all experimental groups.
